# Altered axonal initial segment development links circuit and K_v_7 dysfunction in an *Fmr1* knockout rat

**DOI:** 10.64898/2026.08.18.745439

**Authors:** Jack F. Webster, Marie Pronot, Michael A. Cousin

**Affiliations:** Institute for Neuroscience and Cardiovascular Research, George Square, University of Edinburgh, Edinburgh, United Kingdom; Simons Initiative for the Developing Brain, George Square, University of Edinburgh, Edinburgh, United Kingdom; Muir Maxwell Epilepsy Centre, George Square, University of Edinburgh, Edinburgh, United Kingdom

## Abstract

Fragile X syndrome (FXS) is the leading monogenetic cause of intellectual disability and autism, yet how basic hippocampal circuit properties evolve across development for this condition remains unclear. Here, we studied CA1 pyramidal neurons in male *Fmr1* knockout rats at postnatal day (P)12–15 and 6–10 weeks using *ex vivo* electrophysiology, pharmacology, imaging and biochemistry. P12–15 knockout neurons showed impaired sustained firing, progressive action potential broadening and enhanced activity-dependent synaptic vesicle replenishment. These defects recovered by 6–10 weeks. These phenotypes were linked to K_v_7 channel dysfunction, as K_v_7 activation altered action potential dynamics and neurotransmission in WT but not *Fmr1* KO neurons. Rather than directly altering K_v_7 channel function, *Fmr1* knockout altered axon initial segment (AIS) development such that K_v_7 became functionally inert. These findings suggest that this drives early, transient K_v_7-dependent CA1 dysfunction in FXS and highlight how alterations in AIS development can have profound functional impacts.

## Introduction

Fragile X syndrome (FXS) is a neurodevelopmental disorder (NDD) characterised by intellectual disability and behavioural deficits including anxiety, attention-deficit hyperactivity disorders and autism-associated behaviours such as social withdrawal and avoidance of eye contact^1^. Pathologically, FXS is caused by a trinucleotide repeat expansion of the CGG sequence within the *FMR1* gene^2–4^. This mutation results in the loss of its gene product Fragile X Messenger Ribonucleoprotein (FMRP), an RNA-binding protein which acts primarily as a translational repressor controlling the synthesis of specific proteins primarily at the postsynapse^2^. However, FMRP also has a presynaptic role, where it controls the gating, activation and density of a cohort of ion channels^5^. These channels include Slack potassium channels^6^, large conductance voltage and calcium-gated big potassium (BK) channels^7,8^ and Ca_v_2.2 calcium channels^9^. The absence of FMRP-dependent regulation of these channels results in action potential broadening, circuit hyperexcitability and altered short-term plasticity^7,10–12^. Furthermore, these phenotypes can be corrected by agonism of various potassium channels, including BK and K_v_7 channels^7,8,13^.

A hallmark feature of NDDs is that phenotypical impairments are highly prominent during the early postnatal ‘critical period’ for neurodevelopment, thus altering the temporal trajectory of brain development during the critical developmental window^14^. This is a feature which is well recapitulated in rodent models of FXS with defects in synaptic plasticity^15,16^ and dendritic spine density^17,18^ observed early in development in *Fmr1* knockout (KO) mice. Furthermore, the ‘GABA switch’, the postnatal timepoint at which GABA signalling changes from a net depolarising effect to hyperpolarisation, is delayed in these mice^19^. Finally, we recently revealed that *Fmr1* KO rat hippocampal neurons display a depression in the presynaptic endocytosis mode activity-dependent bulk endocytosis (ADBE)^20^; which appears to recover with culture maturity^21^. Hence, there is now a growing body of literature that shows that *Fmr1* KO rodent models display phenotypical defects at both the pre-and postsynapse that are prominent early in development. However, whether these synaptic defects are sustained over the course of development in *Fmr1* model systems remains unclear.

In this study, we examined these traits in intact circuits from *Fmr1* KO rats at two key timepoints which model the developmental trajectory across both the critical period (Postnatal day 12-15; henceforth referred to as P12-15), and after circuits are established (6-10 weeks old). Implementing an approach of *ex vivo* electrophysiology, immunohistochemistry and protein biochemistry, we explored defects in neuronal firing patterns, synaptic transmission, neurotransmitter release probability and synaptic vesicle (SV) pool dynamics. We found that *Fmr1* KO rats exhibited defects in both action potential discharge and half-width, in addition to enhanced neurotransmission during the early postnatal period. However, these defects were transient in nature, and fully recovered by the time rats reached adolescence. Furthermore, we revealed that these early developmental defects are linked to K_v_7 potassium channels, since the impact of channel-specific agonists were occluded at *Fmr1* KO synapses, most likely due to their redistribution within a remodelled axonal initial segment (AIS).

## Results

### Intrinsic neuronal properties and baseline neurotransmission are unaffected throughout development in *Fmr1* KO rats in the hippocampal CA1 region

To begin to determine which dysfunctional synaptic phenotypes are retained through development, we first assessed a specific subset of parameters in both WT and *Fmr1* KO rats. This was achieved via *ex vivo* patch-clamp electrophysiology of CA1 pyramidal neurons in acute slices from the dorsal hippocampus of both WT and *Fmr1* KO rats at two developmental timepoints: P12-15 which models the critical period of human brain development^22–24^, and P42-70 (6-10 weeks) after circuits are proposed to be established ^22,23,25^.

We first characterised intrinsic neuronal properties (Tables S1 & S2 for full characterisation datasets). In line with previous work in our model^26^ we focussed on resting membrane potential (RMP), input resistance and rheobase (the lowest current step injected which induces action potential firing) as an initial phenotype screen. At P12-15, we observed a non-significant trend towards a more hyperpolarised RMP in neurons recorded from *Fmr1* KO rats (Fig. 1Ai). Both input resistance (Fig. 1Aii) and rheobase (Fig. 1Aiii) were unchanged. In 6-10 week-old rats, all intrinsic neuronal properties in neurons recorded from *Fmr1* KO rats were comparable to WT littermates (Fig. 1Bi-iii). Furthermore, all properties of the rheobase action potential were unchanged at both developmental timepoints (Tables S1 & S2). Therefore, intrinsic neuronal properties within the CA1 hippocampal region were unaffected in *Fmr1* KO rats at both developmental timepoints.

**Figure 1:**
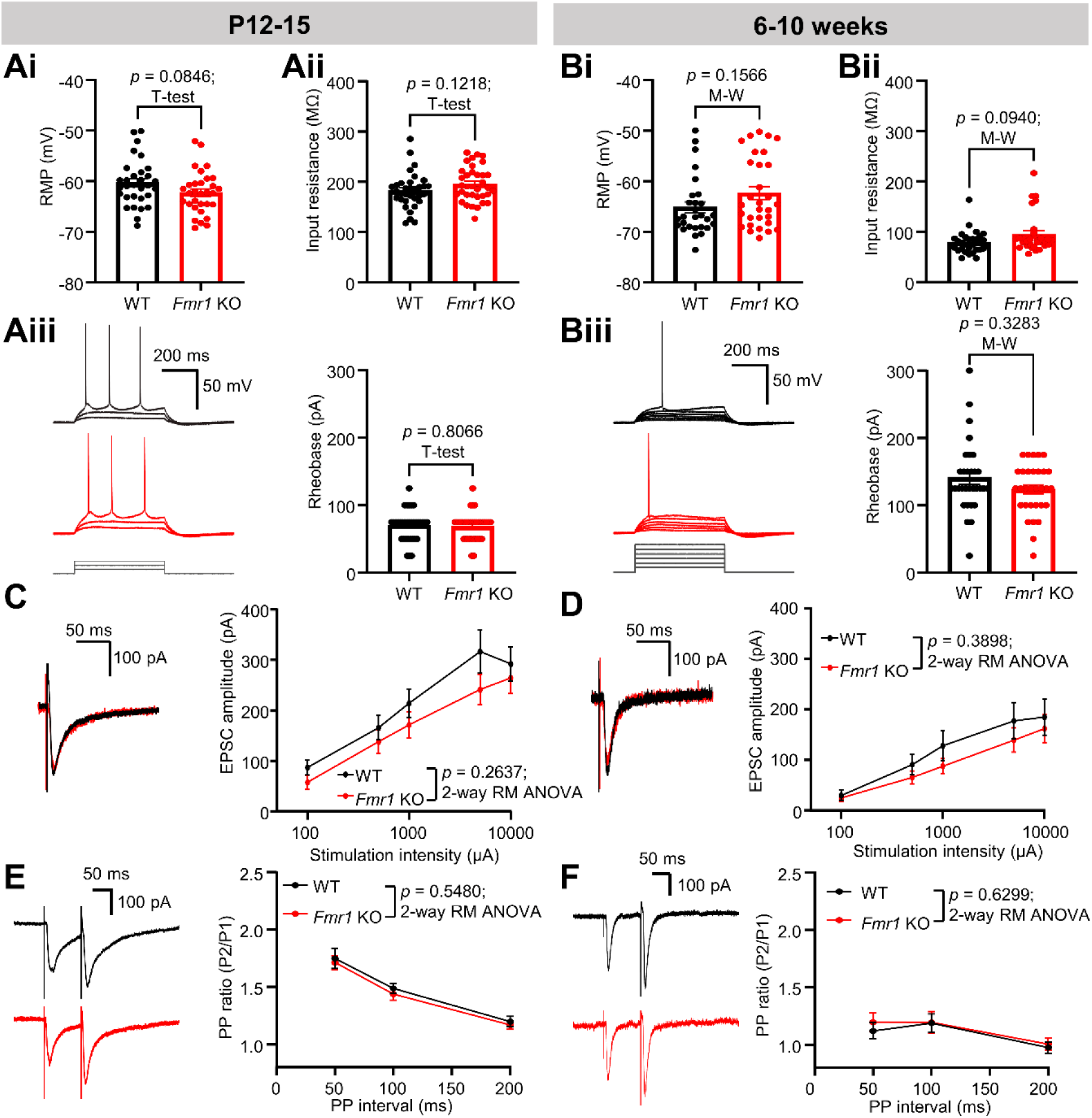
Intrinsic properties and basic neurotransmission in male *Fmr1* KO rats in CA1 hippocampus are unaffected throughout development. **(A)** Comparison plots of intrinsic properties of CA1 pyramidal neurons recorded in acute slices from P12-15 *Fmr1* KO vs WT rats. (Ai) Resting Membrane potential (31/11 WT vs 31/13 KO neurons / rats); (Aii) input resistance (33/11 WT vs 33/13 KO) and (Aiii) rheobase (34/11 WT vs 35/13 KO neurons / rats). Example traces shown on left have current steps of 25 pA increments. (**C**) Left: Example trace of an EPSC elicited in a CA1 pyramidal neuron following presynaptic SC stimulation and; right: input-output plot of EPSC amplitudes across increasing stimulation intensity in slices from P12-15 *Fmr1* KO vs WT rats. 34/11 WT vs 35/13 KO neurons / rats. Example trace = 10000 µA stimulation. (**E**) Left: example traces of recordings in CA1 pyramidal neurons following presynaptic SC paired pulse stimulation and right; plot of paired pulse ratio across stimulation interval in P12-15 *Fmr1* KO vs WT rats. 32/11 WT vs 33/13 KO neurons / rats. (**B**, **D** and **F**) As for (**A**, **C** and **E**); in 6–10-week old rats. (**B**) RMP = 28/11 WT vs 30/8 KO neurons / rats; input resistance = 31/11 WT vs 31/8 KO; rheobase = 34/11 WT vs 35/8 neurons / rats. (**D**) 26/11 WT vs 21/8 KO neurons / rats. (**F**) 21/11 vs 20/8 KO. All summary data are mean ± SEM.

We recently demonstrated a number of SV recycling and endocytosis deficits which occur in primary neuronal cultures derived from *Fmr1* KO rats^20^. The impact of such deficits can be captured in acute slices as defects in baseline neurotransmission^27,28^ and thus we next determined whether this was affected at synapses onto CA1 pyramidal neurons. To achieve this, we utilised electrical stimulation of the presynaptic afferent Schaffer Collateral (SC) pathway, which synapses onto CA1 pyramidal neurons. First, we determined the extent of neurotransmission evoked via stimulation with a single 200 µs pulse of increasing intensity. In slices from both P12-15 and 6–10-week-old rats, we revealed that evoked excitatory postsynaptic current (EPSC) amplitudes were unchanged when genotypes were compared (Fig. 1C & D). To determine whether the intrinsic fusogenicity of SVs were altered at the presynapse in our two independent developmental timepoints, we employed a paired-pulse stimulation protocol to assess presynaptic release probability. This was also unchanged between *Fmr1* KO animals and WT controls at both developmental time points (Fig. 1E & F). Finally, we assessed spontaneous synaptic input, since cultured hippocampal neurons from *Fmr1* KO rats display an increase in spontaneous EPSC frequency, but not miniature EPSC frequency^29^. When sEPSC frequency and amplitude was assessed, both P12-15 (Fig. S1A & B) and 6–10-week-old rats (Fig. S1C & D) displayed no genotype-dependent differences. Thus, both intrinsic neuronal properties and baseline neurotransmission were unaltered by knockout of the *Fmr1* gene at either developmental timepoint in our rat model.

### *Fmr1* KO alters action potential discharge and replenishment of SV pools in P12-15, but not 6-10-week-old rats

As we observed no genotype differences in basic intrinsic neuronal properties or baseline neurotransmission evoked via a single stimulus, we next assessed whether genotype-dependent differences could be revealed during more intense neuronal activity such as during continuous action potential firing. Previous work in P15-25 mice revealed that *Fmr1* KO promoted a pronounced deficit in action potential discharge patterns in hippocampal pyramidal neurons. This induced a robust broadening of the action potential during continuous stimulation, which blocked the capacity of neurons to sustain firing^7^. As such, we next tested whether this phenotype is conserved in P12-15 rats. A depolarising current injection into CA1 pyramidal neurons in *Fmr1* KO rat slices resulted in a strong trend towards a reduction in action potential discharge count when compared to WT controls (Fig. 2A), suggesting an inability to sustain neuronal firing in *Fmr1* KO rats. This was corroborated by a pronounced increase in the action potential half-width in these rats (Fig. 2C), which manifested as an increasing broadening of action potentials during the evoked train (Fig. 2C). Strikingly, this phenotype was absent in slices from 6–10-week-old rats, with CA1 pyramidal neurons recorded from *Fmr1* KO rats discharging action potentials with similar frequencies to WT littermates (Fig. 2B), which did not broaden during increasing depolarising current injection (Fig. 2D). Therefore, the loss of capacity to sustain action potential firing in *Fmr1* KO rats is transient in nature and only present during early development.

**Figure 2:**
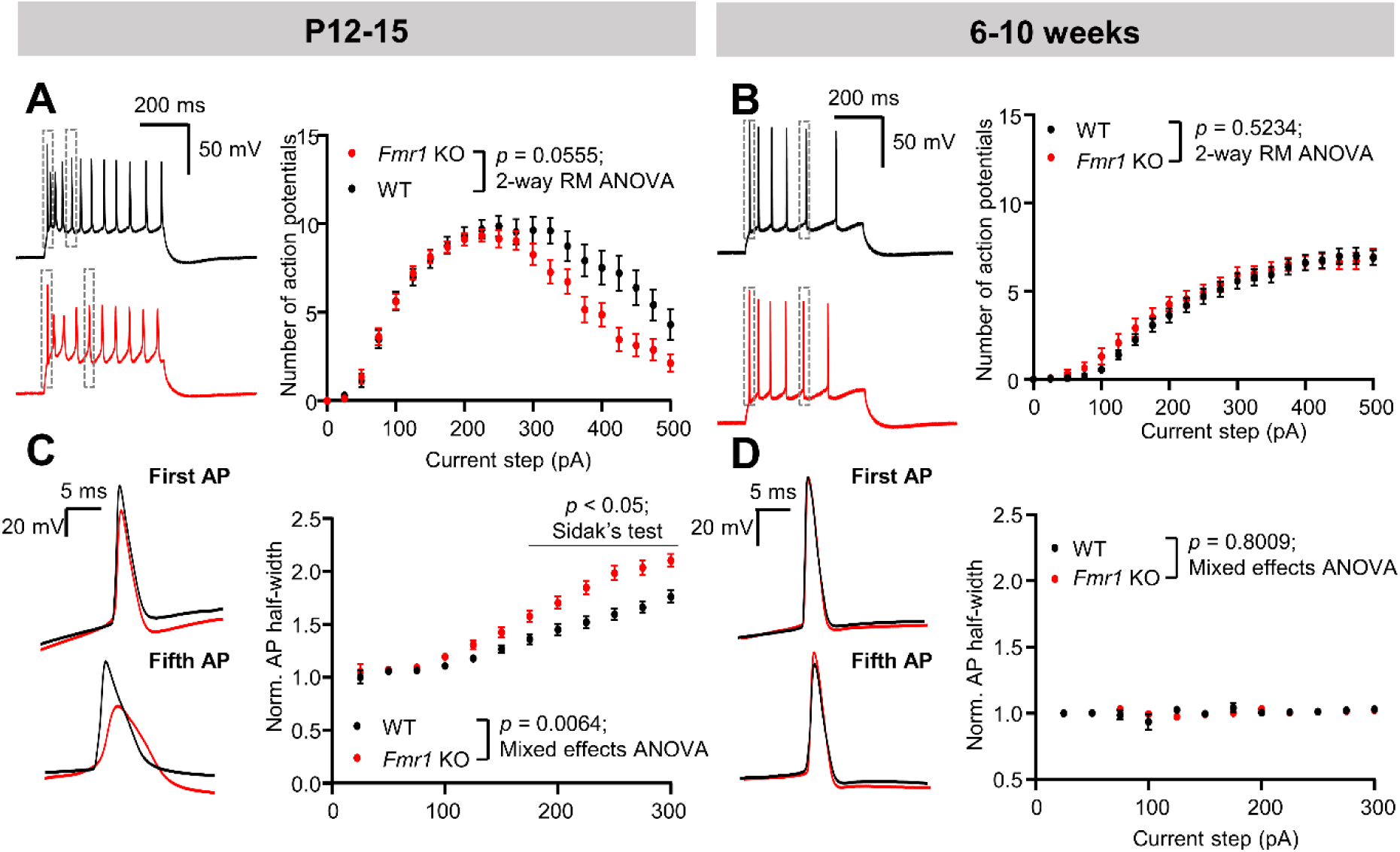
*Fmr1* KO alters action potential discharge in P12-15, but not 6-10-week-old rats: (**A**) Left: example traces of action potential trains recorded in CA1 pyramidal neurons during a depolarising current step injection (300 pA amplitude; 500 ms duration) and; right: input-output curve of action potential firing in response in to depolarising current steps of increasing amplitude in P12-15 *Fmr1* KO vs WT rats. 34/11 WT vs 35/13 KO neurons / rats. (**C**) Left: example traces illustrating action potential broadening in the first vs fifth action potential elicited during current injection in both conditions. Examples are insets of action potentials outlined in (**A**). Note that the half-width of each action potential increases in *Fmr1* KO relative to WT littermates along the duration of each current step. Right: action potential half-width normalised to the mean of the first WT value across increasing depolarising current step injections in both conditions. 34/11 WT vs 35/13 KO. (**B** and **D**) as for (**A** and **C**), in 6–10-week-old rats. (**B**) 31/11 WT vs 31/8 KO neurons / rats. (**D**) 29/11 WT vs 27/8 KO neurons / rats. All data are mean ± SEM.

The loss of capacity to sustain action potential firing in P12-15 *Fmr1* KO rats may lead to alterations in neurotransmitter release, since action potential broadening may result in enhanced presynaptic calcium influx and consequent alterations in the extent of SV fusion. In support, previous studies have reported that *Fmr1* KO increases the size of the SV recycling pool^10,20^; which is defined as the number of SVs available to be accessed via action potential stimulation^30^. However, whether this enhancement is sustained through synaptic maturation remains unknown. Thus, we next sought to assess how neurotransmission is sustained during action potential trains in both P12-15 and 6-10-week-old rats. Using SC pathway stimulation as above (Fig. 1C-F), we assessed current transfer during a 40 Hz stimulus train of 600 action potentials (Fig. 3A & B), which provides a readout of both the capacity to sustain neurotransmission during intense activity, and the replenishment rates of SV pools^27,28^.

**Figure 3:**
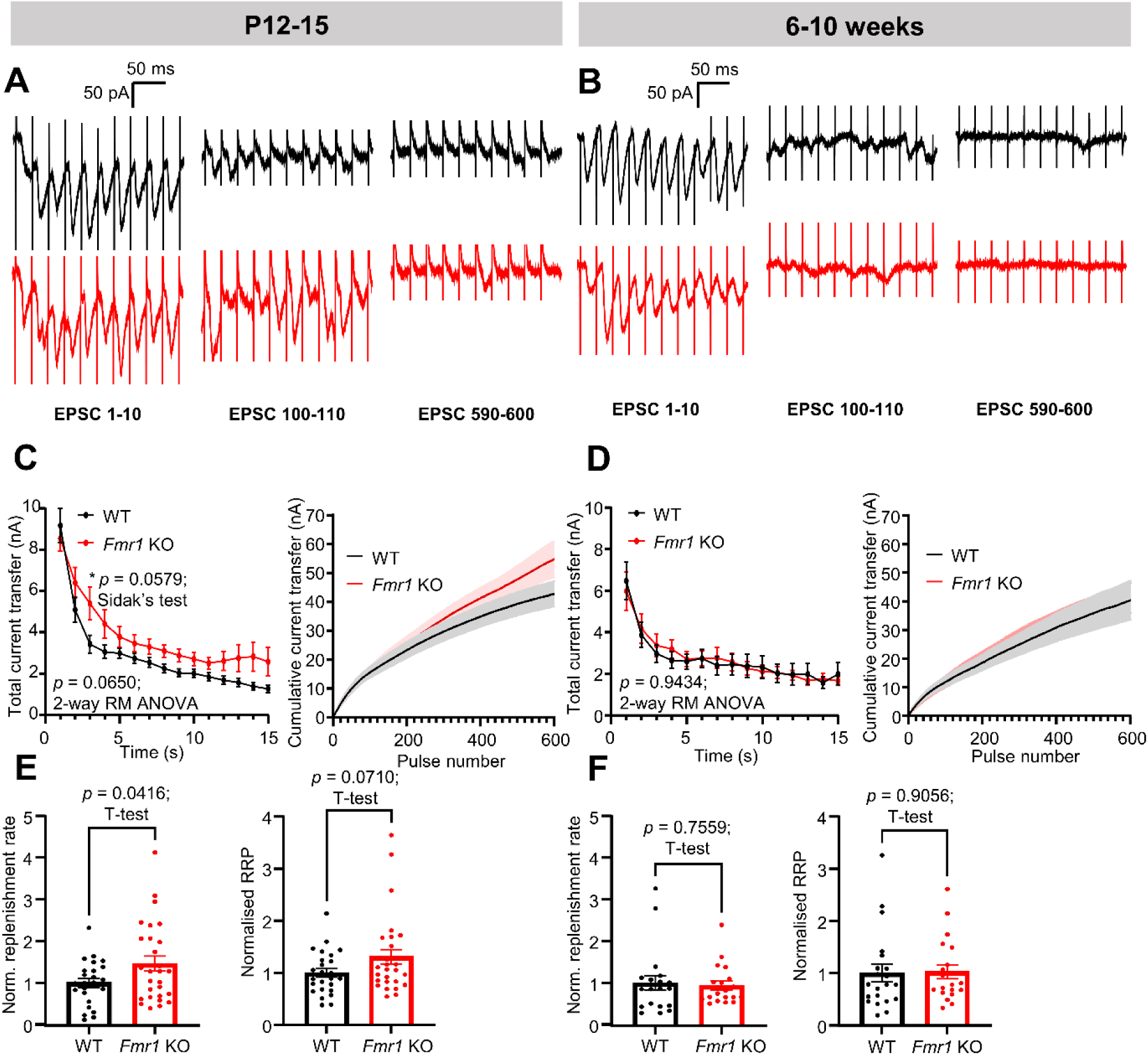
*Fmr1* KO enhances SV pool replenishment in P12-15, but not 6-10-week-old rats: **(A)** Example traces recorded in CA1 pyramidal neurons from *Fmr1* KO vs WT rats during a presynaptic SC stimulation train of 600 action potentials at 40 Hz showing clusters of ten EPSCs recorded at the beginning, after 2.5 seconds, and the end of the train. Note that the EPSC amplitude is slightly larger, specifically during the 2.5 second EPSC cluster in *Fmr1* KO rats. (**C**) Left: Total current transfer recorded in CA1 pyramidal neurons during the 40 Hz presynaptic stimulation protocol outlined in (**A**) binned as total current per second and right: cumulative current transfer over the course of stimulation in P12-15 *Fmr1* KO vs WT rats. 25/11 WT vs 29/13 KO neurons / rats. (**E**) Left: comparison of SV replenishment rates and right: RRP estimates in P12-15 *Fmr1* KO vs WT rats, normalised to the mean of WT controls. 25/11 WT vs 29/13 KO neurons / rats for both comparisons. (**B, D** and **F**) As for (**A, C** and **E**), in 6–10-week-old rats. (**D**) 21/10 WT vs 20/8 KO neurons / rats. (**F**) 21/10 WT vs 20/8 KO neurons / rats for both comparisons. All summary data are mean ± SEM.

In P12-15 *Fmr1* KO rats, we observed a strong trend towards an increase in total current transfer over the course of stimulation, equating to increased neurotransmission (Fig. 3C; *p* = 0.0650; two-way repeated measures ANOVA), with enhanced current transfer most prominent early in the stimulation protocol (Fig. 3A & C). This enhancement was corroborated by a greater SV replenishment rate (Fig. 3E) and a trend towards an increased RRP size in *Fmr1* KO synapses compared to WT (Fig, 3E). Thus, there appears to be an enhancement in the replenishment of the recycling SV pool in *Fmr1* KO animals.

To differentiate between dynamic replenishment of the recycling SV pool and an increase in the overall size of the static RRP, we employed the cyclin-dependent kinase 5 inhibitor roscovitine, which redistributes SVs from the action potential-inaccessible resting pool to the recycling pool^31^ (Fig. S2A). The rationale is that roscovitine will increase the size of the RRP in both WT and *Fmr1* KO synapses, removing the requirement for activity-dependent replenishment of the wider SV pool. Interestingly, roscovitine abolished all apparent genotype-dependent defects in total current transfer (Fig. S2B), replenishment rate (Fig. S2B) and RRP size (Fig. S2B). Therefore, this data suggests that the observed defects are indicative of an increase in replenishment of the SV recycling pool in *Fmr1* KO synapses, as opposed to a general increase in the size of the static RRP.

We next determined whether this enhanced replenishment of the recycling SV pool in *Fmr1* KO rats was retained in 6-10-week-old animals. Importantly, total current transfer (Fig. 3D), SV replenishment rate (Fig. 3F) and RRP size (Fig. 3F) were comparable between WT and *Fmr1* KO rats. Furthermore, application of roscovitine had no impact on these parameters (Fig. S2C). Therefore, the altered SV replenishment observed in *Fmr1* KO rats is transient and recovers by adolescence. Thus, our data collectively reveal that two presynaptic defects in *Fmr1* KO rats, namely the capacity to sustain action potential firing and replenishment of the recycling SV pool, are temporally regulated and only occur during a timeframe equivalent to the critical period for brain development.

### CA1 phenotypical defects in P12-15 *Fmr1* KO rats are linked to loss of K_v_7 channel function

Having revealed two developmentally regulated alterations in presynaptic function in the hippocampus in *Fmr1* KO rats, we next sought to determine the mechanism responsible for these alterations in action potential broadening and increased SV pool replenishment. From this point onwards we focussed exclusively on P12-15 rats, due to the apparent lack of phenotype in their 6–10-week-old counterparts. Recent work has shown that K_v_7 potassium channel dysfunction mediates some of the neuronal excitability defects within the hippocampus in *Fmr1* KO mice^13^. Importantly, these channels are widely acknowledged to be dysfunctional in many NDD’s^32^. Since neuronal excitability defects in a FXS mouse model have also been linked to alterations in presynaptic plasticity^10^, we next explored the hypothesis that altered K_v_7 channel function may underlie the defects in action potential firing (Fig. 2) and SV pool replenishment (Fig. 3).

To achieve this, we repeated the experiments above in slices from P12-15 *Fmr1* KO rats and WT littermates treated with either the K_v_7 channel activator retigabine, or the vehicle DMSO. Pre-treatment with retigabine (2.5 µM) did not significantly alter passive intrinsic properties nor the properties of the rheobase action potential in WT (Table S3) or *Fmr1* KO (Table S4) neurons. However, as expected, retigabine robustly reduced neuronal excitability in CA1 pyramidal neurons recorded in slices from WT rats, both increasing the rheobase current step (Fig. 4A) and reducing the number of action potentials fired during depolarising current step injection (Fig. 4B). Furthermore, retigabine pre-treatment reduced the action potential half-width in slices from WT rats (Fig. 4C), in agreement with a reduction in excitability^7^. Remarkably, the capacity of retigabine to reduce excitability was almost entirely lost in CA1 neurons recorded from *Fmr1* KO rats. Specifically, rheobase in slices from *Fmr1* KO animals was unchanged in comparison to DMSO controls (Fig. 4A) and the number of action potentials fired during depolarising current injection was also comparable overall (Fig. 4B), although post-hoc multiple comparisons did reveal statistically significant differences specifically between 75-125 pA current steps. This loss-of-effect was also reflected in the action potential half-width, which was unchanged in *Fmr1* KO neurons relative to vehicle controls following retigabine treatment (Fig. 4C).

**Figure 4:**
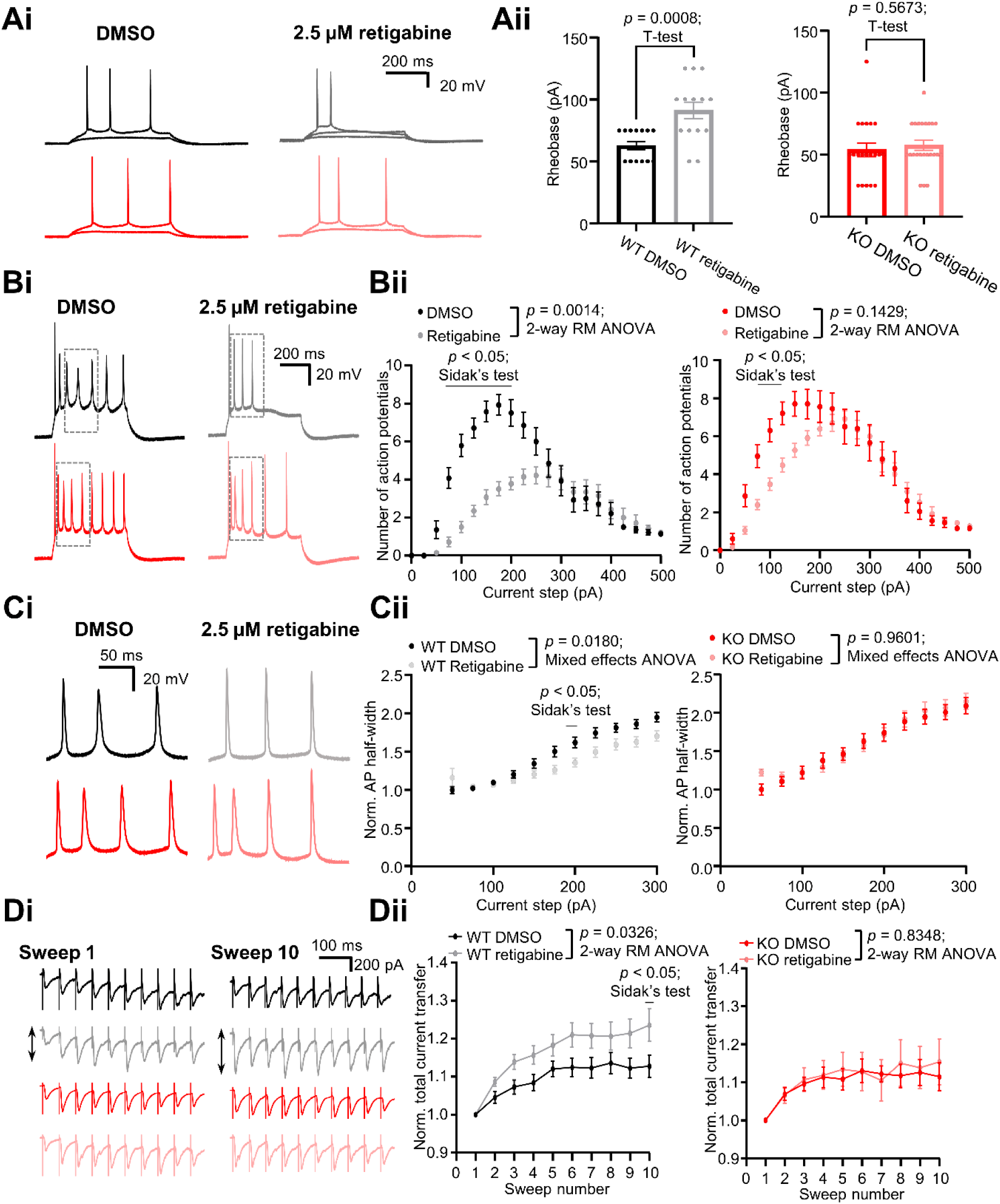
Defects in action potential discharge and neurotransmission in P12-15 *Fmr1* KO rats are linked to loss of K_v_7 channel function. (**Ai**) Example traces of rheobase recordings in CA1 pyramidal neurons from P12-15 *Fmr1* KO vs WT, pretreated with either 0.1% DMSO or 2.5 µM retigabine. Note the increase in rheobase following retigabine pretreatment specifically in WT neurons. Example traces = 175 pA current injection. (**Aii**) Comparison plots of neuronal rheobase in CA1 pyramidal neurons recorded from; left: WT rats treated with either 2.5 µM or 0.1% DMSO (14/5 DMSO vs 14/5 retigabine neurons / rats) or; right: the same conditions in *Fmr1* KO rats (20/9 DMSO vs 23/9 retigabine neurons / rats). (**Bi**) Example traces illustrating action potential firing during depolarising current injection recorded in under the same conditions outlined in (**A**). (**Bii**) Input-output curve of action potential firing across increasing depolarising current steps in conditions outlined in (**A**). WT = 14/5 DMSO vs 14/5 retigabine; KO = 20/9 DMSO vs 23/9 retigabine neurons / rats. (**Ci**) Example traces illustrating action potential broadening during depolarising current injection recorded in CA pyramidal neurons from *Fmr1* KO vs WT rats treated as above. Example traces are zooms of the boxed regions in (**Bi**). Note that retigabine causes narrowing of the action potential only in WT neurons. (**Cii**) Action potential half-width normalised to the first DMSO value across increasing depolarising current step injections in above conditions. WT = 42/18 DMSO vs 40/18 retigabine; KO = 32/16 DMSO vs 39/16 retigabine neurons / rats. (**Di**) Example traces illustrating EPSC trains elicited during a presynaptic 20 Hz SC stimulation train of 10 action potentials recorded in conditions as above. For each condition, 10 sweeps in total were recorded with a 10 s delay between sweeps. Example traces show the first and last sweep under each condition. Note that retigabine causes facilitation of neurotransmission only in WT neurons (arrows). (**Dii**) Total current transfer recorded across each sweep under the stimulation conditions outlined in (**Di**), normalised to sweep 1 of the DMSO control for each genotype. WT = 27/13 DMSO vs 25/13 retigabine; KO = 12/7 DMSO vs 15/7 retigabine neurons / rats. All summary data are mean ± SEM.

We next determined if alterations in K_v_7 channel function at *Fmr1* KO synapses could contribute to the enhancement of SV pool replenishment we observed (Fig. 3). We tested this by repeating the protocol of a tetanic 40 Hz afferent stimulation train of presynaptic SCs while recording in postsynaptic CA1 neurons following pre-treatment with retigabine. Retigabine treatment resulted in a statistically significant increase in total current transfer in WT slices, specifically during the initial phase of the action potential train (Fig. S3Ai). However, the total current transfer over the duration of the stimulus was comparable between DMSO-and retigabine-treated WT slices (Fig. S3Ai) and *Fmr1* KO slices (Fig. S3Aii). In addition, extrapolated values for both SV replenishment and RRP size were comparable following retigabine treatment in both WT (Fig. S3Bi) and KO slices (Fig. S3Bii).

The apparent decoupling of the impact of K_v_7 modulation on action potential dynamics from SV dynamics in *Fmr1* KO rats suggests that they are driven via divergent mechanisms. However, the observed alteration in SV fusion by retigabine during the first second of the stimulation train in WT slices (Fig. S3Ai) indicates that the influence of K_v_7 channels on neurotransmitter release may only be prominent during less intense stimulation. Thus, we tested the effect of retigabine in WT and *Fmr1* KO slices during a stimulation train of 10 action potentials at 20 Hz repeated with a 10 s delay between sweeps. This protocol recruits SVs to replenish the recycling pool, and acts as an indicator of presynaptic short-term plasticity^33^. We quantified total current transfer during each sweep by measuring the area under the curve of each EPSC as a proxy for cumulative neurotransmitter release and observed a significant enhancement by retigabine in WT slices (Fig. 4D). Strikingly, retigabine was unable to facilitate neurotransmission to any observable extent in *Fmr1* KO slices (Fig. 4D). Thus, K_v_7 channels do appear to be capable of modulating replenishment of the SV recycling pool, but only during lower intensity stimulation. Critically, this capacity is absent in *Fmr1* KO synapses, consistent with occlusion of the impact of retigabine on action potential dynamics.

Interestingly, this *Fmr1* KO-specific K_v_7 channel loss-of-function was not observed during K_v_7 blockade (Fig. 5). For example, pre-treatment with the K_v_7 channel blocker XE991 (5 µM) shifted the action potential discharge curve to the left in CA1 pyramidal neurons recorded from both WT and *Fmr1* KO slices, indicative of an increase in excitability. Although there was not a statistically significant effect of genotype alone in either WT or KO slices (Fig. 5A & B), the 2-way ANOVA genotype x current step interaction statistic was significantly different in both (WT *p* = 0.0014; KO *p* < 0.0001) reflecting the altered shape of the curve in both genotypes following XE991 pre-treatment. XE991 pre-treatment also induced a similar broadening of the action potential half-width in both WT (Fig. 5C) and *Fmr1* KO slices (Fig. 5D). XE991 did however appear to have mild, yet statistically significant, genotype-specific effect on RMP and action potential threshold (Tables S5 & S6). However, it did not have an observable effect on either SV pool replenishment or RRP size in either WT or *Fmr1* KO slices (Fig. S3C & D).

**Figure 5:**
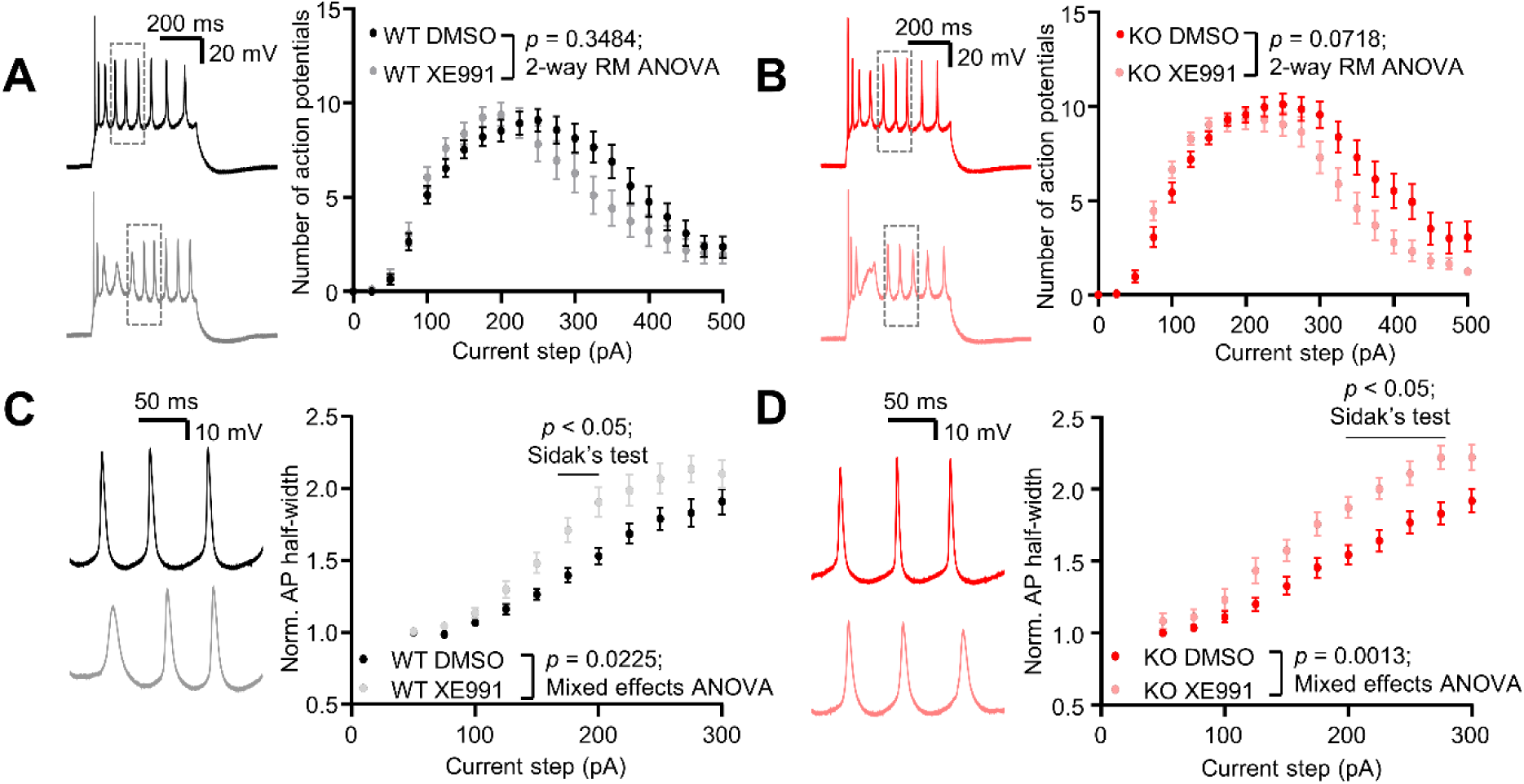
K_v_7 blockade broadens action potentials in both P12-15 *Fmr1* KO and WT rats. (**A**) Left: example traces of action potential trains recorded in CA1 pyramidal neurons during a depolarising current step injection (300 pA amplitude; 500 ms duration) and; right: input-output curve of action potential firing in response to depolarising current steps of increasing amplitude in P12-15 WT rats treated with either 0.1% DMSO or 5 µM XE991. 25/10 DMSO vs 22/9 XE991 neurons / rats. (**C**) Left: example traces illustrating action potential half-width during current injection in both conditions. Examples are insets of action potentials outlined in (**A**). Note that XE991 pretreatment causes action potential broadening. Right: action potential half-width normalised to the mean of the first DMSO value across increasing depolarising current step injections in both conditions. 25/10 DMSO vs 22/9 XE991 neurons / rats. (**B** and **D**) as for (**A** and **C**), in *Fmr1* KO rats. (**B**) 28/10 DMSO vs 25/10 XE991 neurons / rats. (**D**) 28/10 DMSO vs 25/10 XE991 neurons / rats. All data are mean ± SEM.

In summary, the combined conclusion from these experiments is that loss of K_v_7 function is likely at least a contributing factor to the action potential dynamic (Fig. 2) and neurotransmission (Fig. 3) phenotypes in our model, as K_v_7 activation fails to modulate these characteristics specifically in the KO’s. However, the non-genotype dependent enhancement of action potential half-width via K_v_7 blockade suggests that these channels are still at least partially active.

### Changes in AIS morphology underpin alterations in K_v_7.2 distribution in *Fmr1* KO hippocampal neurons

Taken together, the above results indicate that K_v_7 channels in the CA1 region of *Fmr1* KO rats exhibit a pronounced alteration of function. However, this does not appear to be due to changes in expression, since their inhibition has comparable effects in both WT and *Fmr1* KO synapses (Fig. 5). Indeed, previous work has reported no observable changes in gross expression of either K_v_7.2 or K_v_7.3 in *Fmr1* KO mice^13^. Because of this, we speculated that the observed phenotypical defects could be mediated by a non-canonical function of FMRP^34^, such as a direct interaction of FMRP with K_v_7.2. In support, direct binding of FMRP to both BK channels and Ca_v_2.2 channels can impact either channel function or trafficking to the plasma membrane respectively^9,35^. We tested this hypothesis using a co-immunoprecipitation assay from P14 rat brain lysates (Fig. S4). Antibodies against FMRP robustly extracted the protein from these lysates in contrast to negative control IgG. However, subsequent Western blotting revealed the absence of K_v_7.2 in FMRP immunoprecipitates (Fig. S4). Therefore, there was no obvious direct interaction between FMRP and K_v_7.2 in rat brain.

As there was no apparent direct interaction between these two proteins and no reported changes in gross K_v_7 hippocampal expression^13^, we next hypothesised that the mechanism underpinning the observed loss of K_v_7 channel function could be linked to alterations in channel clustering specifically along *Fmr1* KO axons. This is because their axonal expression is known to powerfully modulate action potential threshold and neuronal RMP in CA1^36^. To test this, we immunostained for K_v_7.2 channels in embryonic hippocampal cultures from *Fmr1* KO and WT littermates at 14 DIV (Fig. S5A & D). This specific point in development was chosen since *Fmr1* KO neurons in culture display similar developmentally transient defects associated with our current observations in whole circuits at this timepoint^21^. K_v_7.2 clustering was assessed using coefficient of variation (CV) analysis^37,38^. This analysis quantifies how variable the distribution of K_v_7.2 channels is along the axon, with a high value reflecting their restriction into clusters, and lower values representing a more diffuse distribution. CV analysis revealed that K_v_7.2 channel clustering was unaltered in *Fmr1* KO cultures compared to control (Fig. S5B & C). Furthermore, gross axonal K_v_7.2 expression was also unchanged when genotypes were compared (Fig. S5B). We also analysed K_v_7.2 expression within other neuronal compartments; namely the dendrites (Fig. S5D-G) and the soma (Fig. S6). Although there was an apparent reduction in K_v_7.2 expression in the dendrites of cultured neurons (Fig. S5E), we could not recapitulate this in intact circuits from fixed brain slices (Fig. S5G). The dendritic CV was unaltered in both conditions (Fig. S5E & G). Furthermore, automated detection of K_v_7.2-positive soma (Methods & Fig. S6A) throughout dorsal CA1 and the entire hippocampal formation revealed no differences in expression patterns between genotypes (Fig. S6B). Therefore, an alteration in gross neuronal K_v_7.2 expression does not underlie the phenotypical defects in action potential broadening and SV replenishment that we report in *Fmr1* KO rats.

The AIS is a known hotspot for K_v_7.2 clustering, where its influence on action potential generation and propagation is effectively amplified relative to the rest of the axon^36,39^. We thus hypothesised that AIS-specific K_v_7.2 alterations may underlie the apparent loss of function in *Fmr1* KO rats (Fig. 4). To test this, we stained fixed brain slices (Fig. 6A) and primary neuronal cultures (Fig. 6E) with the AIS marker Ankyrin-G to allow morphological quantification of the AIS in both WT and *Fmr1* KO neurons. In brain slices, we observed a reduction in the overall density of labelled AIS within the CA1 pyramidal layer of *Fmr1* KO rats (Fig. 6A), which was accompanied by a trend towards a reduction in Ankyrin-G fluorescence signal along the course of the AIS (Fig. 6B-D). However, we did not detect any obvious genotype-dependent differences in AIS morphology in fixed tissue (Fig. 6B). In primary neuronal cultures, Ankyrin-G staining revealed that while the total AIS area was unaltered in *Fmr1* KO neurons (Fig. 6F), AIS length was shorter (Fig. 6F) and its width was broader (Fig. 6F) in neurons from *Fmr1* KO rats. This results in an overall “stubbier” AIS in *Fmr1* KO neurons (Fig. 6E). Gross K_v_7.2 expression at the AIS was unaltered in *Fmr1* KO primary neuronal cultures (Fig. 6G & H), as was its CV (Fig. 6H), consistent with the absence of genotype-dependent channel expression (Fig. S5A-C) in axons. We could not reliably quantify K_v_7.2 expression at the AIS in fixed tissue, primarily due to poor antibody penetration.

**Figure 6:**
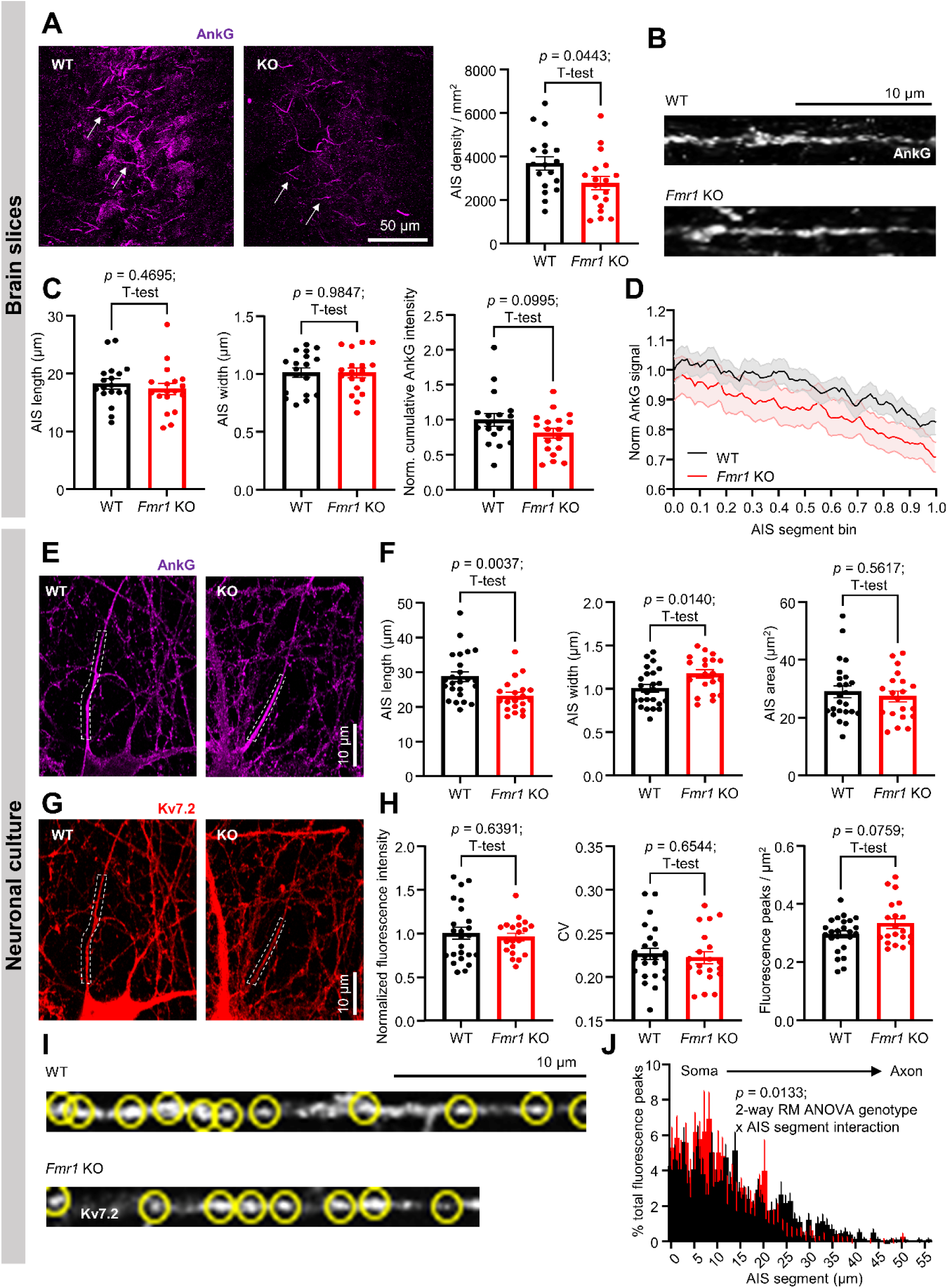
Developmental dysregulation of the axonal initial segment underpins alterations in K_v_7.2 distribution in *Fmr1* KO hippocampal neurons. **(A)** Left: example images depicting Ankyrin-G staining profiles in WT vs *Fmr1* KO fixed hippocampal brain slices. Arrows indicate labelled AIS. Right; comparison of labelled AIS density between conditions. (**B**) Representative zoom images of the AIS in WT vs *Fmr1* KO brain slices. (**C**) Comparison plots of AIS length, width and total cumulative Ankyrin-G staining intensity between conditions. (**D**) Relative Ankyrin-G staining intensity binned to normalised AIS segment in WT vs *Fmr1* KO brain slices. 18/6 WT vs 18/6 KO slices / rats. (**E**) Example images depicting Ankyrin-G staining profiles in WT vs *Fmr1* KO primary hippocampal cultures. The AIS is denoted by the broken-lined box. Note the shortened and wider shape of the AIS in the *Fmr1* KO example relative to WT. (**F**) Comparison plots quantifying morphological characteristics of the AIS in WT vs *Fmr1* KO neurons (length, width and area). 24/6 WT vs 20/6 KO coverslips / embryos for all comparisons. (**G**) Example images depicting K_v_7.2 staining profiles from the same neurons in (**E**). (**H**) Comparison plots of normalised fluorescence intensity, coefficient of variability (CV), and detected fluorescence peaks / µm^2^ in the AIS of WT vs *Fmr1* KO neurons. 24/6 WT vs 20/6 KO coverslips / embryos for all comparisons. (**I**) Example images of background-corrected K_v_7.2 intensity profiles derived from the AIS segments denoted in (**G**). Yellow circles indicate fluorescence peaks. (**J**) Frequency-distribution histogram illustrating differences in K_v_7.2 fluorescence peak distribution in WT vs *Fmr1* KO neurons indicating relative peak densities at AIS segments relative to distance from soma. Data are coverslip means of total % peaks detected binned to 1 µm intervals ± SEM. 24/6 WT vs 20/6 KO coverslips / embryos.

We also quantified K_v_7.2 fluorescence peak maxima in primary neuronal cultures, which is an indication of high fluorescence signal relative to the surrounding area. Using this analysis, we observed a trend towards an increased density of fluorescence peaks in *Fmr1* KO neurons (Fig. 6H). We speculated that this effect may be a direct result of the observed alterations in AIS morphology in cultured neurons (Fig. 6G & H) thus resulting in a concentration of K_v_7.2 expression nearer the soma in *Fmr1* KO neurons. Indeed, fluorescence peak distribution analysis confirmed this hypothesis, with *Fmr1* KO neurons displaying an overall altered distribution of fluorescence peaks, with higher density relative to WT neurons proximal to the soma, but progressively decreasing with distance (Fig. 6I & J). Taken together, this fixed tissue and neuronal culture data suggest that gross K_v_7.2 expression is unaltered in the axon of *Fmr1* KO rats. However, alterations in the AIS likely underpin its loss of function. Previous developmental studies of the AIS have revealed that the AIS rapidly lengthens^40–42^ and Ankyrin-G labelling density increases^41^ during the early postnatal period in rodents. Our data are thus consistent with perturbed development of the AIS in *Fmr1* KO neurons relative to WT neurons, which may ultimately serve to limit K_v_7.2 control of action potential generation and propagation, thus explaining its profound reduction of influence in *Fmr1* KO rats (Fig. 4).

### K_v_7 channels are functionally intact in *Fmr1* KO neurons

Our above data indicate a profound loss of capacity of K_v_7 channels to modulate action potential dynamics in immature *Fmr1* KO neurons (Fig. 4), even though the expression of these channels appears broadly unaffected. Thus, an open question is whether these channels are still functionally intact. To test this, we recorded the M-current, the slow hyperpolarising potassium current conducted via these channels, in both WT and *Fmr1* KO CA1 pyramidal neurons (Fig. 7). This current could be clearly isolated pharmacologically in neurons from both genotypes (Fig. 7A), with M-current amplitude displaying no genotype-dependent difference (Fig. 7Aii). Furthermore, 2.5 µM retigabine robustly increased M-current amplitude in both genotypes (Fig. 7B). Thus, despite their loss of capacity to modulate action potential dynamics, K_v_7 channels appear functionally intact in immature *Fmr1* KO CA1 pyramidal neurons.

**Figure 7:**
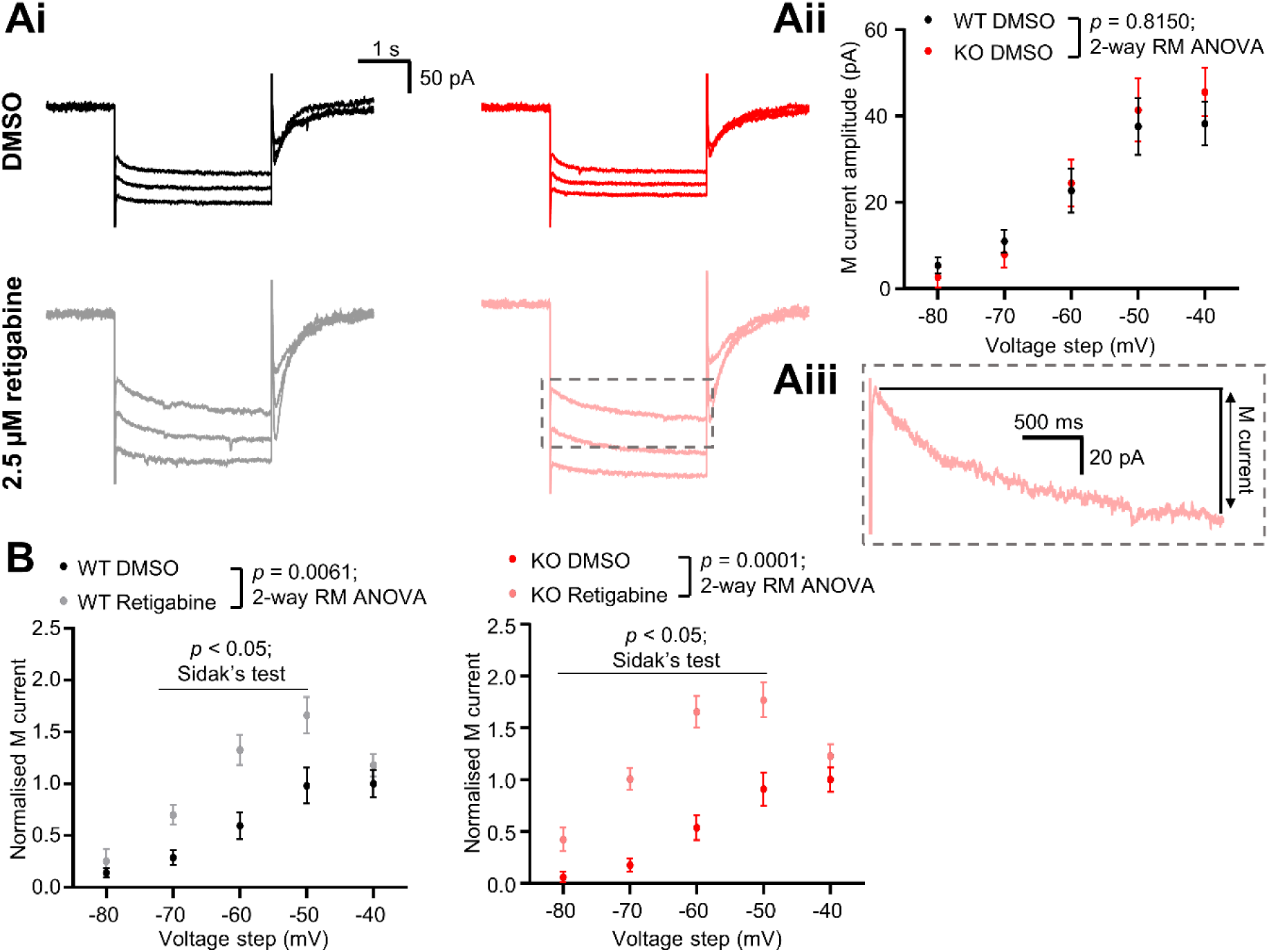
K_v_7 channels are functionally intact in P12-15 *Fmr1* KO rats. (**Ai)** Example current traces showing M-current induced following hyperpolarising voltage steps in WT (left) and *Fmr1* KO (right) CA1 pyramidal neurons treated with either 0.1% DMSO (top row) or 2.5 µM retigabine (bottom row). Example traces are from-50 to-70 mV voltage steps. (**Aii**) Current-voltage plot comparing M-current elicited across voltage steps in WT vs *Fmr1* KO neurons. WT = 19/6 DMSO vs 18/6 retigabine; KO 13/5 DMSO vs 13/5 retigabine neurons / rats. Data are mean ± SEM. (**Aiii**) Inset zoom of the boxed region in (Ai). M-current is calculated as the difference between the peak and steady state current over the duration of the voltage injection. (**B**) Current-voltage plots from both WT (left) and *Fmr1* KO (right) CA1 pyramidal neurons comparing the fold-increase in M-current amplitude following 2.5 µM retigabine preincubation relative to 0.1% DMSO preincubation. WT = 19/6 DMSO vs 18/6 retigabine; KO 13/5 DMSO vs 13/5 retigabine neurons / rats. Data are mean ± SEM normalised to the mean of the-40 mV step for the DMSO condition.

## Discussion

The accurate establishment of neuronal circuits in early life is critical for typical brain development. However, it is still unclear as to what aspects of synaptic dysfunction during this critical period are maintained to adulthood, and which are corrected in later life. In this study we investigated whether phenotypical defects within the hippocampal CA1 region in the *Fmr1* KO rat model of human FXS were retained after this critical period had completed. For this, we chose two different age groups which are known to correspond to the human developmental timepoints of infancy^22,23^ and adolescence^22,23^. A key finding is that the action potential broadening and enhanced SV pool replenishment observed in P12-15 *Fmr1* KO rats are strikingly transient, since both fully recover by adolescence. Furthermore, this developmentally regulated phenotype appears to be linked to defects in K_v_7 channel function: a group of potassium channels which mediate the M-current and defects in development of the AIS; both of which have previously been linked to neurodevelopmental disorders^32,43^.

Our findings of synaptic dysfunction in P12-15 *Fmr1* KO rats closely correlate with previous studies in *Fmr1* KO mice at the same developmental timepoint, with both action potential broadening and presynaptic plasticity defects in hippocampal slices observed in this model system^7,10^. However, our critical finding is that these phenotypes are fully corrected in later life. The mechanism of this recovery of function is still undetermined, however a parsimonious explanation is that *Fmr1* KO neurons exhibit a delay in maturation, reaching full maturity at a later timepoint. In support, synaptic dysfunction has been reported in a series of FXS models 2-3 weeks after birth, with these parameters recovering with maturity^15–19,21^. Furthermore, previous work has revealed that the action potential half-width progressively narrows during early development^44,45^ thus implying that the alterations in action potential broadening and firing observed in P12-15 *Fmr1* KO rats relative to WT controls is indicative of developmental delay. We acknowledge that while our data highlight alterations in AIS development as a possible mechanism to explain the transient phenotypical defects we observe, we have not tested if this phenotype is resolved by adolescence. Previous work in our *Fmr1* KO rat model has assessed AIS length in two to three-month old *Fmr1* KO rats and found this to be comparable to WT controls^26^. Thus, we assume this phenotype is also transient and likely normalised by adolescence.

We also acknowledge that initially, our findings that *Fmr1* KO neurons appear to fire less action potentials than WT neurons during intense current injection (Fig. 2A) appears in contradiction to the prominent circuit hyperexcitability theory of FXS^34^. Our explanation is that the prominent action potential broadening (Fig. 2C) exhibited by *Fmr1* KO neurons result in a firing modality where the kinetics of action potential discharge are too slow to support sustained firing, thus causing the neurons to enter a state of depolarization block where they fire a reduced number of action potentials. Depolarization block is a known consequence of excessive neuronal activity^46^ and thus we believe our data are therefore consistent with circuit-level hyperexcitability.

In the wider context of the literature, action potential broadening has been observed previously in CA3 neurons from P15-25 *Fmr1* KO mice^7^ However in this instance this phenotype was driven by inhibition of BK channels, rather than loss of K_v_7 function as our data suggest (Fig. 4). In contrast, we report no obvious differences in neuronal afterhyperpolarisation in our rat model (Tables S1 and S2), a neuronal property mediated by BK channel function^47^. This implies possible mechanistic differences between the rat and mouse models; specifically, we speculate that the influence of BK channels within the rat may be less significant than in the mouse. Conversely in rat, K_v_7 channels appear capable of modulating action potential half-width, a function which is lost in *Fmr1* KO rats (Fig. 4). To our knowledge, this has not been reported in mouse. These key findings implicate potassium channel function as a possible area of species divergence, which likely has implications for both the study of NDDs and fundamental neurobiology. ^7^

As stated above, defects in action potential discharge count and half-width in our *Fmr1* KO rat model are at least partially mediated by dysfunction in K_v_7 channels (Fig. 4 and 5). However, a key apparent paradox is that the capacity of K_v_7 channels to modulate the action potential is largely occluded in *Fmr1* KO CA1 neurons (Fig. 4); yet these channels are apparently unaltered in terms of gross expression (Figs. 6, S5 and S6), are fully functional (Fig. 7) and susceptible to blockade (Fig. 5). This pattern is consistent with altered functional coupling; i.e. the channels are still present, but are no longer positioned such that increasing their open probability produces a strong physiological effect.

Our AIS staining data (Fig. 6) provide an anatomical substrate for such decoupling. As discussed above, our data are consistent with developmental alterations in the AIS^40–42^ in *Fmr1* KO neurons. These alterations effectively result in an altered distribution of K_v_7.2 early in neurodevelopment (Fig. 6). Because K_v_7 channels exert disproportionate control over excitability when concentrated in the AIS^36,39^, even subtle shifts in the precise expression pattern of K_v_7 channels within the AIS can strongly alter their impact on action potential initiation and propagation. In this model, AIS remodelling in the *Fmr1* KO reduces K_v_7 leverage over the spike-generating compartment. Indeed, it is known that alterations in AIS length serve as an excitability tuning mechanism^48,49^. Specifically, a longer AIS presents a greater distance for AIS-anchored sodium and potassium conductance to exert their influence over spike kinetics, thus resulting in a greater effect^49^. Following this logic, a shorter AIS would permit a lesser influence of such ionic conductance, entirely consistent with our data. This therefore, offers an explanation as to why retigabine no longer narrows spikes or enhances transmission, despite XE991 still having an observable effect on spike broadening in both genotypes (Fig. 5). Thus, a key takeaway from our work is that even apparently minor changes in AIS development can have profound functional implications for ion channels anchored to the AIS; even if the channels themselves are not altered in any observable way.

A highly notable observation is that while our results are consistent with a recent study using an *Fmr1* KO mouse model^13^ in that K_v_7 channel dysfunction is a prominent feature of both the rat and mouse models, the underlying pathology appears to substantially differ. Specifically, in the *Fmr1* KO mouse, retigabine treatment is sufficient to reduce the neuronal hyperexcitability observed in this model to WT levels^13^. This implies that K_v_7 dysfunction may directly contribute to neuronal hyperexcitability in this model, whereas retigabine exhibited a pronounced loss-of-effect in *Fmr1* KO rats relative to WT controls (Fig. 4). This apparent disparity may also be explained by differences in AIS morphology. In contrast to our rat data, *Fmr1* KO mice exhibit a longer AIS within CA1^50^, which would be consistent with an enhanced K_v_7.2 conductance^49^. Thus, while K_v_7 channel dysfunction may be a core mechanism of rodent models of FXS, it is possible that the underlying mechanism may differ in the mouse, as these channels still appear physiologically functional in this model. Interestingly, a recent study showed that pathological mutations in the *KNCQ2* gene, which encodes K_v_7.2, was sufficient to induce dysfunction consistent with delayed neural network development^51^ in neurons differentiated from human induced pluripotent stem cells. This suggests that K_v_7 channel dysfunction could be causative in the apparent developmental delay of our model, and perhaps neurodevelopmental disorders in general. Thus, correction of K_v_7 channel dysfunction may offer a valid therapeutic target to normalise neurodevelopmental trajectory in NDD’s.

## Conclusions

Our work highlights the developmentally transient nature of phenotypical abnormalities in the CA1 region of the hippocampus in a rat model of FXS, and the profound impact that alterations in AIS development can have on ion channel function. This work is thus consistent with the premise of delayed circuit development in NDD’s^14^, and the potential of a critical therapeutic window to maximise the efficacy of therapeutic interventions. We do note that while our study focuses on FXS, differing NDD’s likely display different developmental time courses, and thus further work in other rodent models of NDD’s should serve to shed light on this. Finally, while we highlight K_v_7 channel dysfunction as a conserved feature in rodent models of FXS, we note that the underlying mechanisms appear to differ between species and thus caution should be exerted when translating these findings to the human condition.

## Materials and methods

### Animals

In all cases, heterozygous *Fmr1^em1/PWC^* female rats^52^ were mated with male wild-type (WT) rats to produce male WT or homozygous *Fmr1^em1/PWC^* rat (referred to as *Fmr1* KO rats throughout this work) littermates. All rats were maintained on a Long-Evans hooded background. Rats were group housed under standard conditions on a 12:12 hour light/dark cycle, at 21 ± 2 °C room temperature with *ad libitum* access to food and water in mixed genotype cages of up to 5 rats. For the preparation of hippocampal cultures, heterozygous female *Fmr1* KO rats and WT males were timed-mated to produce embryos which were used for cultures at embryonic day 18. Rat genotype information was retained by an individual not involved in the study, and released to the investigators when required.

All procedures were performed in accordance with the UK Animal (Scientific Procedures) Act 1986, under Project and Personal licence authority and were approved by the Animal Welfare and Ethical Review Body at the University of Edinburgh (Home Office Project Licence - PP5745138 and PP6320781).

### Acute brain slice preparation

For acute brain slice preparation, animals were first rendered unconscious by exposure to isoflurane (Covetrus) anaesthesia and then intraperitoneally administered with a terminal overdose of sodium pentobarbital (brand name Dolethal, Vetoquinol). Once animals were no longer responsive to a noxious toe-pinch stimulus, they were transcardially perfused with a solution of ice-cold sucrose-based artificial cerebrospinal fluid (sACSF) bubbled with carbogen (95% O_2_; 5% CO_2_). sACSF consisted of (in mM) sucrose 75, NaCl 87, KCl 2.5, NaHCO_3_ 25, NaH_2_PO_4_ 1.25, MgCl_2_ 7, CaCl_2_ 0.5, glucose 25. Animals were perfused for approximately 90 seconds a rate of 20 ml / minute after which animals were decapitated, and brains were rapidly removed and transferred to a beaker of ice-cold carbogenated sACSF. Brains were then sectioned on a Leica VT1200 vibratome (Leica Biosystems). Sections containing the dorsal hippocampus, the structure in which all electrophysiology and fixed tissue experiments were performed, were cut at 350 µm intervals in the coronal plane, and then immediately transferred to a beaker of carbogenated sACSF warmed to 37 °C where they remained for 30 minutes. Slices were then transferred to a beaker of room temperature standard ACSF which contained (in mM) NaCl 125, KCl 2.5, NaHCO_3_ 25, NaH_2_PO_4_ 1.25, MgCl_2_ 1, CaCl_2_ 2 and glucose 10 where they remained until required for electrophysiology. Details of all reagents used in acute slice preparation and *ex vivo* electrophysiology can be found in table 1.

**Table 1:** Details of all electrophysiological reagents.

| Reagent | Supplier | Catalogue no. | CAS no. |
| --- | --- | --- | --- |
| Sucrose | Sigma-Aldrich | S8501 | 57-50-1 |
| NaCl | Sigma-Aldrich | S9888 | 7647-14-5 |
| KCl | Sigma-Aldrich | P3911 | 7447-40-7 |
| NaHCO<sub>3</sub> | ThermoFisher | C12336.A1 | 144-55-8 |
| NaH<sub>2</sub>PO<sub>4</sub> | ThermoFisher | 447760010 | 13472-35-0 |
| CaCl<sub>2</sub> | BioWorld | 40320005 | 10043-52-4 |
| MgCl<sub>2</sub> | Honeywell | 63020 | 7786-30-3 |
| Glucose | Sigma-Aldrich | G8270 | 50-99-7 |
| K-Gluc | Sigma-Aldrich | P1847 | 299-27-4 |
| EGTA | Sigma-Aldrich | E3889 | 67-42-5 |
| HEPES | ThermoFisher | BP310 | 7365-45-9 |
| Na<sub>2</sub>ATP | Merck Millipore | A7699 | 34369-07-8 |
| NaGTP | Merck Millipore | G8877 | 36051-31-7 |
| Na<sub>2</sub>-phosphocreatine | Merck Millipore | P7936 | 19333-65-4 |
| Biocytin | Merck Millipore | B4261 | 576-19-2 |

### *Ex vivo* electrophysiology and analysis

Acute slices were transferred to a slice bath on a Scientifica SliceScope Pro 1000 electrophysiology rig and perfused continuously with sACSF at a flow rate of 2-3 ml / minute. For pathway stimulation experiments, a bipolar stimulation electrode made of 0.05 mm thick twisted Nichrome (80% nickel / 20% chromium; Advent Research Materials, NI6530) wire connected to a DS3 constant-current stimulation box (Digitimer) was implanted into *stratum radiatum* of the proximal CA1 region to stimulate the Schaffer collateral pathway. Patch-pipettes were pulled on a Sutter P-97 Flaming / Brown type micropipette horizontal puller (Sutter Instruments) from glass capillaries of dimensions as follows: inner diameter 0.86 mm, outer diameter 1.5 mm, length 100 mm (Harvard Apparatus, G1510F-4) of typical tip resistance of 3-4 MΩ and were filled with a potassium gluconate (K-gluc) based internal solution which consisted of (in mM) K-gluc 142, KCl 4, EGTA 0.5, HEPES 10, MgCl_2_ 2 Na_2_ATP 2, NaGTP 0.3, Na_2_-phosphocreatine 1, biocytin 2.7.

For patch-clamp experiments, the CA1 pyramidal cell layer was visualised under Differential Interference Contrast microscopy with a SciCam Pro digital camera (Scientifica) using a 40x water-immersion objective lens. CA1 pyramidal neurons were approached and a seal was formed via application of negative pressure. Short negative pressure pulses were then applied until whole-cell configuration was achieved, with access resistance typically <15 MΩ. Access resistance was monitored throughout, and recordings were discarded if it changed by more than 20%.

Recordings were made using a Multiclamp 700B amplifier (Molecular Devices) and filtered using a Digidata 1550B digitizer (Molecular Devices). All recordings were acquired in Clampex version 10.0 (Molecular Devices) at a sampling rate of 20 kHz. For intrinsic property recordings, input resistance was assessed in voltage clamp by acquiring a-10 mV pulse (duration 500 ms) while neurons were clamped at-70 mV. All other intrinsic properties were acquired in current clamp. Firing properties were assessed via injection of a series of current steps (-125 – 500 pA; 25 pA increments; 500 ms duration) and sag potential was acquired via injection of a-300 pA step (500 ms duration). Synaptic neurotransmission data were acquired as previously described^27,28,53^. Briefly, neurons were voltage-clamped at-80 mV and EPSCs were evoked using single stimuli of increasing intensity ranging from 100 – 10000 µA (200 µs duration). The stimulation intensity was then adjusted to an amplitude to evoke an EPSC of ≈ 200 pA. Neurotransmitter release probability was assessed using a paired pulse stimulation protocol (50 – 200 ms pulse interval; 3 sweeps per interval; 30 s delay between sweeps). SV dynamics and pool size were assessed using two separate protocols: a 20 Hz train of 10 action potentials, repeated over 10 sweeps (10 s delay between sweeps), and a high-intensity 40 Hz stimulation of 600 action potentials performed for 1 sweep. For experiments requiring drugs, slices were preincubated with 100 μM roscovotine (APExBio, A1723), 2.5 μM retigabine (APExBIO, B5888) or 5 μM XE-991 (Sigma-Aldrich, X2254) for 15 minutes before being transferred to the recording chamber, where experimentation was carried out as above.

M-current experiments were performed in sACSF supplemented with 100 µM 4-aminopyridine (Merck-Millipore, 275875), 500 nM tetrodotoxin (Hellobio, HB1035) and 10 µM ZD7288 (Abcam, AB120102) as described previously^54^. Whole-cell configuration was achieved and neurons were voltage clamped at-30 mV. Hyperpolarizing voltage steps (-10 to-50 mV; 10 mV increments; 3 s duration) were used to elicit the M current. As above, slices were preincubated with either 0.1% DMSO or 2.5 µM retigabine.

For electrophysiology data analysis, action potential count following depolarising current injection was calculated using the AP count feature in Stimfit 0.15.8. Most other measurements including intrinsic property, EPSC and M-current measurements were carried out in Axograph X 1.8.0, using the event detection feature to analyse action potential amplitude and kinetics, and the measure peaks and shapes feature to measure EPSC or M-current amplitude and kinetics. Unless otherwise specified, all intrinsic properties were derived from the rheobase action potential (i.e. the first action potential elicited during depolarising current injection). For 20 Hz and 40 Hz stimulation train experiments, raw trace sample points were exported to a Microsoft excel spreadsheet and EPSCs amplitudes were calculated using custom peak detection macros. Total current transfer (the area under the curve) was calculated using a custom script written in R Studio. SV mobility estimates RRP size and SV replenishment rates) were calculated as previously described^27,28^. Briefly, the cumulative current transfer was calculated for each EPSC during a 40 Hz stimulation train of 15 seconds duration (see *ex vivo* electrophysiology section), and a linear regression was fit to the values for the last 1 second of stimulation. The y-intercept and gradient were utilised as the RRP and SV replenishment rate estimates, respectively.

### Primary neuronal cultures

Pregnant female rats were killed by exposure to increasing CO_2_ concentration (starting low and rising until death), followed by cervical dislocation to confirm death. Embryos were removed from the pregnant female and then killed by decapitation. Hippocampi were individually dissected from male embryos and then digested in a 10 µl/mL papain solution (Worthington Biochemical, LK003178) at 37°C for 20 minutes. Papain was then neutralised using DMEM/F12 (ThermoFisher Scientific, 21331-020) supplemented with 10% fetal bovine serum (BioSera, S1810-500) and 1% penicillin / streptomycin (ThermoFisher Scientific, 15140-122). Cells were triturated to form a single cell suspension and plated at 5 x 10^4^ cells per coverslip on laminin (10 µg/ mL; Sigma-Aldrich, L2020) and poly-D-lysine (Sigma-Aldrich, P7886) coated 25 mm glass coverslips (VWR International Ltd). The cultures were maintained in Neurobasal media (ThermoFisher Scientific, 21103-049) supplemented with 2% B-27 (ThermoFisher Scientific, 17504-044), 1% penicillin / streptomycin and 0.5 mM L-glutamine (ThermoFisher Scientific, 25030-024). At day *in vitro* (DIV) 2-3, 1 μM cytosine arabinofuranoside (Sigma-Aldrich, C1768) was added to the media to inhibit glial proliferation.

### Perfusion fixation and cryosectioning

For perfusion fixation, P12-15 rats were first terminally anaesthetised and transcardial perfusion was performed as described above, with the exception that the animals were sequentially perfused with ice-cold phosphate buffered saline (PBS, Sigma-Aldrich, P4417) and then 4% paraformaldehyde (w/v; Agar Scientific, AGR1018) in PBS. Brains were then rapidly dissected and post-fixed in 4% paraformaldehyde in PBS overnight. This solution was then exchanged with a cryoprotective solution of 30% sucrose in PBS (w/v) and the brains maintained until they dropped to the bottom of the tube. At this point, the solution was replaced and renewed, and once the brain dropped to the bottom a second time, the brain was considered ready to section. For cryosectioning, brains were embedded in optimal cutting temperature solution (Agar Scientific, AGR1180), frozen and sectioned at 50 µm on a freezing microtome (Leica Biosystems). Sections were stored in PBS at 2-8 °C until required for immunohistochemistry.

### Immunohistochemistry

Coverslips containing primary neuronal cultures were fixed for 10 minutes with a solution of 4% paraformaldehyde in PBS. Coverslips were then washed twice with PBS and blocked for 1 hour in a solution of 5% fetal bovine serum (BioSera, S1810-500) and 0.1% Triton X-100 (Sigma-Aldrich, T8787) in PBS at 4 °C. Coverslips were then incubated overnight with a primary antibody cocktail consisting of rabbit anti-K_v_7.2 (1/1000 dilution), mouse anti-Neurofilament (1/200 dilution) and 0.1% Triton X-100 in PBS at 4 °C; or guinea pig anti-Ankyrin-G (1/500 dilution). Coverslips were then washed in PBS and incubated with a secondary antibody cocktail consisting of goat anti-mouse conjugated to Alexa fluor 488 (1:500 dilution) and donkey anti-rabbit conjugated to Alexa fluor 647 (1:500 dilution); or goat anti-guinea pig conjugated to Alexa fluor 568 (1:500 dilution), in PBS for 1 hour at room temperature in darkness. Coverslips were then washed repeatedly in PBS, washed once in deionised water, and mounted on microscope slides (ThermoFisher Scientific, BS7011/2) using Fluoromount mounting media with DAPI added to stain nuclei (Electron Microscopy Sciences, 17984-24). Full details of all antibodies used within this study can be found in Table 2.

**Table 2:** Details of all antibodies.

| Antibodies | Supplier | Cat no. | RRID |
| --- | --- | --- | --- |
| mouse anti-FMR1 | Sigma-Aldrich | SAB4200597 | AB_3695994 |
| mouse anti-IgG | BD Biosciences | 550875 | AB_393942 |
| mouse anti-neurofilament | BioLegend | 801901 | AB_2566782 |
| rabbit anti-Kv7.2 | Synaptic Systems | 368103 | AB_2620130 |
| guinea pig anti-Ankyrin G | Synaptic Systems | 386 005 | AB_2737033 |
| goat anti-mouse, Alexa 488 | ThermoFisher | A11017 | AB_2534084 |
| goat anti-guinea pig, Alexa 488 | ThermoFisher | A11073 | AB_2534117 |
| goat anti-guinea pig, Alexa 568 | ThermoFisher | A11075 | AB_2534119 |
| donkey anti-rabbit, Alexa 647 | ThermoFisher | A31573 | AB_2536183 |
| anti-Mouse IRDye 800CW | LI-COR Biosciences | 925-32212 | AB_2716622 |
| anti-Rabbit IRDye 680RD | LI-COR Biosciences | 925-68073 | AB_2716687 |

Brain section immunohistochemistry was performed with sections free-floating in the relevant media. Sections were washed twice in PBS and blocked in a solution of 5% fetal bovine serum and 0.3% Triton in PBS for 3 hours 4 °C. Primary antibody incubations were performed overnight in a cocktail consisting of rabbit anti-K_v_7.2 (1/250 dilution) and guinea pig anti-Ankyrin-G (1/500 dilution) supplemented with 0.3% Triton. Slices were washed and then incubated in a cocktail of secondary antibodies consisting of donkey anti-rabbit conjugated to Alexa fluor 647 (1:500 dilution) and goat anti-guinea pig conjugated to Alexa fluor 488 (1:500 dilution) at room temperature for 3 hours. Slices were then washed three times with PBS and mounted using Fluoromount mounting media with DAPI,

### Image acquisition and analysis

For imaging axons and dendrites in primary neuronal cultures, coverslips were imaged on a Leica THUNDER widefield imaging system (Leica Microsystems) equipped with a multi-line high-intensity LED fluorescence light source and controlled using LAS X software. Two fluorescence channels were acquired; both with a 63x oil immersion objective (1.40 numerical aperture, NA). Channel 1 (corresponding to the anti-neurofilament stain) was acquired using an L5 filter set (excitation bandpass 480/40, dichroic 505, emission bandpass 527/30) with LED intensity set to 10% and 200 ms exposure to image axons and 30% intensity / 750 ms exposure for dendrites; and channel 2 (corresponding to the anti-K_v_7.2 stain) was acquired using a Y5 filter set (excitation bandpass 620/60, dichroic 660 LP, emission bandpass 700/75) with LED intensity set to 17% and 750 ms exposure time for axons and 17% intensity / 400 ms exposure for dendrites.

Hippocampal overview images of fixed brain slices were also acquired on the Leica THUNDER widefield imaging system using a 20x dry objective, 0.8 NA, and the LAS X tile scan function. Two fluorescence channels were acquired. Channel 1, corresponding to DAPI, was acquired using the Leica A4 UV/DAPI filter set (excitation bandpass 360/40, dichroic 400, emission bandpass 470/40), with LED intensity set to 10% and an exposure time of 100 ms. Channel 2, corresponding to K_v_7.2, was acquired using the Y5 filter set with LED intensity set to 55% and an exposure time of 1 s. Large-volume computational clearing was applied post hoc to the acquired overview images.

For axon initial segment imaging, images were acquired on a Leica TCS SP8 confocal microscope (Leica Microsystems) controlled with LAS X software. Two fluorescence channels were collected using a 63x oil-immersion objective (NA 1.40) with a 1024 × 1024 frame size, no digital zoom, and single optical sections (single plane). Channels were acquired sequentially at 200 Hz with no frame averaging (1 frame) for both channels. Channel 1 (anti-Ankyrin-G) was excited with the 552 nm laser line (1% laser power) and detected with a PMT detector over 593–617 nm; PMT gain was set to 800 V. Channel 2 (anti-K_v_7.2) was excited with the 638 nm laser line (2.5% laser power) and detected with a HyD detector in standard mode over 643–789 nm. Spectral detection/unmixing was not used.

Images of fixed brain slices were acquired using a 63x high-resolution objective on a Leica TCS SP8 as for coverslips above, with parameters adjusted as follows. Channel 1 (anti-AnkyrinG) was excited with the 488 nm laser line (1.5% laser power) and channel 2 (anti-K_v_7.2) was excited with the 638 nm laser line (2% laser power). Both channels were acquired sequentially at 100 Hz with no frame averaging. A confocal z-stack was acquired for each slice consisting of the most superficial 10 µm of the slice (1 µm optical section).

Both epifluorescence and confocal images were analysed in Fiji (ImageJ) using custom macros. Confocal z-stacks were first pre-processed by converting to a maximum intensity z-projection. For analysis of individual neuronal compartments, the axonal or dendritic segment of interest was first defined by manually tracing a segmented-line region of interest along the appropriate neuronal segment (i.e. the axon, AIS or dendrite) and the macro computed segment length as the cumulative arc-length of the traced vertices. The traced segment was then straightened (straighten width = 2 µm for axonal segments; 5 µm for dendrites), and the same straightening transform was applied to the K_v_7.2 channel to enable measurements as a function of distance along the neuronal segment. K_v_7.2 fluorescence was quantified along the straightened segment by extracting an intensity profile along the X-axis (mean intensity across the strip height at each X position). Profiles were exported both in raw, uncorrected form and after background correction using rolling-ball subtraction (radius = 1.5 µm). Spatial heterogeneity of K_v_7.2 expression was analysed using the coefficient of variation (CV = SD/mean) of the background-corrected profile. K_v_7.2 puncta were identified on the background-corrected straightened image using Fiji’s Find Maxima in point mode (bright peaks on dark background; tolerance = 30% of intensity range), with a minimum inter-peak separation (0.25 µm). For analyses of general axonal segments, channel 1 (neurofilament) was used solely to select the axonal portion for analysis and no additional measurements were performed in this channel. For AIS analyses, channel 1 (AnkyrinG) was additionally quantified and used to generate a binary AIS mask by automated thresholding (Otsu with a minimum threshold floor). This mask enabled estimation of AIS width along the segment and constrained K_v_7.2 peak counts to peaks colocalised with the AnkyrinG-defined AIS; K_v_7.2 mean intensity within the AIS mask was also computed (raw and background-corrected).

Automated detection of K_v_7.2-positive soma in hippocampal overview images was performed using a second macro which operated as follows. Two-channel images were split into K_v_7.2 and DAPI channels, and anatomical regions of interest (ROIs) were manually defined on the DAPI channel. Rolling-ball background subtraction and Gaussian blurring were applied to the K_v_7.2 channel before automated detection. K_v_7.2-positive soma were detected as local intensity maxima within each ROI using ImageJ’s Find Maxima function. For each ROI, the macro calculated ROI area, K_v_7.2 mean intensity, K_v_7.2-positive neuron count, density, and mean detected-neuron intensity. A verification TIFF containing ROI boundaries and detected soma was also generated for visual quality control.

### Immunoprecipitation

P14 rats were killed by exposure to an increasing concentration of CO_2_, and after decapitation brains were rapidly removed and were solubilized for 1 h at 4 °C in lysis buffer (10 mM Tris-HCl, 10 mM EDTA, 150 mM NaCl, 1 % Triton X-100, 0.1 % SDS, pH 7.5) supplemented with protease inhibitor cocktail. Lysates were sonicated for 10 s using an MSE Soniprep 150 sonicator and clarified at 15,000×*g* at 4 °C for 15 minutes with a HettichLab Mikro 22R benchtop centrifuge. The resulting detergent-soluble supernatant (400 µg) was incubated in 1.5ml Eppendorf tubes with either 4 µg of mouse anti-FMRP or 4 ug of IgG in lysis buffer overnight at 4 °C with 30 µl of Protein G Sepharose™ 4 Fast Flow Resin. The bead/lysate suspension was precipitated by centrifuging at 250*g* at 4 °C. Supernatant was discarded, with beads resuspended in 1 ml lysis buffer before centrifugation as before. This washing protocol was repeated three times. After the final wash, proteins were eluted by the addition of sodium dodecyl sulphate (SDS) sample buffer (2% SDS, 10 % glycerol, 5 % β-mercaptoethanol, bromophenol blue, 62.5 mM Tris, pH 7.4) and subsequent heating at 60° for 10 minutes.

### Immunoblotting

Protein extracts were resolved on an 8 % SDS-PAGE gel and transferred onto a 0.2 µm nitrocellulose membrane. The membrane was incubated for 1 hour in Odyssey blocking buffer then immunoblotted with mouse anti FMRP (1/1000 dilution) and rabbit anti-K_v_7.2 (1/1000) overnight at 4°C in Odyssey blocking buffer. After five washes in blocking buffer, membranes were incubated with two secondary antibodies from LI-COR: anti-Mouse IRDye 800CW and anti-Rabbit IRDye 680RD in blocking buffer. Membranes were imaged on an Odyssey M Imaging System (LI-COR Biosciences) using LI-COR Image Studio Lite software (version 5.2). Details of all general reagents utilised in immunoprecipitation and blotting can be found below in table 3.

**Table 3:** Details of all immunoprecipitation and blotting reagents.

| Reagent | Supplier | Cat no. | CAS no. |
| --- | --- | --- | --- |
| Tris (Trizma) base | Merck | 93352 | 77-86-1 |
| EDTA | Sigma-Aldrich | E-5134 | 6381-92-6 |
| Sodium Chloride | Fisher | S/3160/63 | 7647-14-5 |
| Triton X-100 | Sigma-Aldrich | T8787 | 9036-19-5 |
| Sodium dodecyl sulfate | Sigma-Aldrich | L5750 | 151-21-3 |
| Protein G Sepharose 4 Fast Flow Resin | Cytiva | 17061801 | N/A |
| EGTA | Sigma-Aldrich | E-4378 | 67-42-5 |
| Glycerol | Fisher | G/0600/17 | 56-81-5 |
| Bromophenol blue | BioRAD | 161-0404 |  |
| β-mercaptoethanol | Sigma-Aldrich | 125472500 | 60-24-2 |
| Protran Premium<br>0.2um NC Membrane | Amersham | 10600004 | N/A |
| 30% Acrylamide/bis | BioRAD | 1610159 | 79-06-1 |
| N,N,N',N'-Tetramethyl-<br>ethylenediamine | Sigma-Aldrich | T7024 | 110-18-9 |
| Ammonium persulfate | Sigma-Aldrich | A-3678-25G | 7727-54-0 |
| Glycine | Sigma-Aldrich | 10070150 | 56-40-6 |
| cOmplete™, EDTA-free Protease Inhibitor<br>Cocktail | Sigma-Aldrich | 04693132001 | N/A |
| Odyssey blocking<br>buffer | LI-COR Biosciences | 927-4000 | N/A |

## Statistics and reproducibility

All data were plotted and statistical analysis performed in GraphPad prism 9.3.1. For pairwise comparisons, a Shapiro-Wilk normality test was performed to assess normality of distribution. Where one or both sets of data were normally distributed, an unpaired t-test (T-test) was used to test for statistical significance. Where both sets of data were not normally distributed, a Mann-Whitney test (M-W) was used. All comparisons were two-tailed, and any *p* value of < 0.05 was considered statistically significant. Repeated measures ANOVA (2-way RM ANOVA) was used to compare datasets where multiple data points were acquired per sample. Mixed effects ANOVA was used for datasets where X-axis datapoints were missing. This was exclusive to the action potential broadening comparisons, as no datapoints could be calculated at current steps where neurons did not fire action potentials. These datasets were analysed up to 300 pA steps of current injection as beyond this point, neurons from P12-15 rats were generally unable to sustain firing in (Fig. 2A) thus resulting in insufficient action potential trains to perform an equitable analysis. For row x column analysis (i.e. ANOVAs), post hoc multiple comparison tests were used to test for statistical significance at each point across the X axis. For statistical comparisons, data are displayed as n/N where n = neurons or coverslips, and N = animals for electrophysiology experiments, or embryos for primary neuronal cultures. n/N values can be found in legends of corresponding figures. In all cases, animal selection was performed by an individual not involved in the study, and the experimenter was kept blind to animal genotype during experimentation.

## Acknowledgements

This work was funded by a grant from the Simons Foundation to MAC (529508) and a Marie-Curie Skłodowska / UK Research and Innovation Guarantee Award to MP (EP/Y024559/1). Drs. James Hogg and Nawon Kim are both thanked for their preparation of primary neuronal cultures.

## Author Contributions

Conceptualization, JFW, MAC; Methodology, MP, JFW; Formal Analysis, MP, JFW; Investigation, MP, JFW, MAC; Resources, MAC; Writing – Original Draft, JFW; Writing – Review & Editing, MP, JFW, MAC; Funding Acquisition, MP, MAC.

## Data availability statement

All data supporting the findings of this study are available within the paper and in the Supplementary Data File. All relevant data can be provided upon reasonable request to the corresponding authors.

## References

1. Cregenzán-Royo, O., Brun-Gasca, C. & Fornieles-Deu, A. Behavior Problems and Social Competence in Fragile X Syndrome: A Systematic Review. Genes (Basel). 13, 280 (2022).

2. Hagerman, R. J., et al. Fragile X syndrome. Nat. Rev. Dis. Primers 3, (2017).

3. Verkerk, A. J. M. H. et al. Identification of a gene (FMR-1) containing a CGG repeat coincident with a breakpoint cluster region exhibiting length variation in fragile X syndrome. Cell 65, 905–914 (1991).

4. Colak, D. et al. Promoter-Bound Trinucleotide Repeat mRNA Drives Epigenetic Silencing in Fragile X Syndrome. Science (1979). 343, 1002–1005 (2014).

5. Ferron, L. Fragile X mental retardation protein controls ion channel expression and activity. J. Physiol. 594, 5861–5867 (2016).

6. Brown, M. R. et al. Fragile X mental retardation protein controls gating of the sodium-activated potassium channel Slack. Nat. Neurosci. 13, 819–821 (2010).

7. Deng, P.-Y. et al. FMRP Regulates Neurotransmitter Release and Synaptic Information Transmission by Modulating Action Potential Duration via BK Channels. Neuron 77, 696–711 (2013).

8. Deng, P. & Klyachko, V. A. Genetic upregulation of BK channel activity normalizes multiple synaptic and circuit defects in a mouse model of fragile X syndrome. J. Physiol. 594, 83–97 (2016).

9. Ferron, L., Nieto-Rostro, M., Cassidy, J. S. & Dolphin, A. C. Fragile X mental retardation protein controls synaptic vesicle exocytosis by modulating N-type calcium channel density. Nat. Commun. 5, 3628 (2014).

10. Deng, P.-Y., Sojka, D. & Klyachko, V. A. Abnormal Presynaptic Short-Term Plasticity and Information Processing in a Mouse Model of Fragile X Syndrome. Journal of Neuroscience 31, 10971–10982 (2011).

11. Wang, X. et al. Activity-dependent regulation of release probability at excitatory hippocampal synapses: a crucial role of fragile <scp>X</scp> mental retardation protein in neurotransmission. European Journal of Neuroscience 39, 1602–1612 (2014).

12. Booker, S. A. et al. Altered dendritic spine function and integration in a mouse model of fragile X syndrome. Nat. Commun. 10, 4813 (2019).

13. Luque, M. A., Morcuende, S., Torres, B. & Herrero, L. Kv7/M channel dysfunction produces hyperexcitability in hippocampal CA1 pyramidal cells of *Fmr1* knockout mice. J. Physiol. 602, 3769–3791 (2024).

14. Meredith, R. M. Sensitive and critical periods during neurotypical and aberrant neurodevelopment: A framework for neurodevelopmental disorders. Neurosci. Biobehav. Rev. 50, 180–188 (2015).

15. Zhang, J., Hou, L., Klann, E. & Nelson, D. L. Altered Hippocampal Synaptic Plasticity in the *Fmr1* Gene Family Knockout Mouse Models. J. Neurophysiol. 101, 2572–2580 (2009).

16. Harlow, E. G. et al. Critical Period Plasticity Is Disrupted in the Barrel Cortex of Fmr1 Knockout Mice. Neuron 65, 385–398 (2010).

17. Nimchinsky, E. A., Oberlander, A. M. & Svoboda, K. Abnormal Development of Dendritic Spines in *FMR1* Knock-Out Mice. The Journal of Neuroscience 21, 5139–5146 (2001).

18. Grossman, A. W. et al. Developmental characteristics of dendritic spines in the dentate gyrus of Fmr1 knockout mice. Brain Res. 1355, 221–227 (2010).

19. He, Q., Nomura, T., Xu, J. & Contractor, A. The Developmental Switch in GABA Polarity Is Delayed in Fragile X Mice. The Journal of Neuroscience 34, 446–450 (2014).

20. Bonnycastle, K., Kind, P. C. & Cousin, M. A. FMRP Sustains Presynaptic Function via Control of Activity-Dependent Bulk Endocytosis. J. Neurosci. 42, 1618–1628 (2022).

21. Kim, N., Bonnycastle, K., Kind, P. C. & Cousin, M. A. Delayed recruitment of activity-dependent bulk endocytosis in *Fmr1* knockout neurons. J. Neurochem. 168, 3019–3033 (2024).

22. Semple, B. D., Blomgren, K., Gimlin, K., Ferriero, D. M. & Noble-Haeusslein, L. J. Brain development in rodents and humans: Identifying benchmarks of maturation and vulnerability to injury across species. Prog. Neurobiol. 106–107, 1–16 (2013).

23. Bell, M. R. Comparing Postnatal Development of Gonadal Hormones and Associated Social Behaviors in Rats, Mice, and Humans. Endocrinology 159, 2596–2613 (2018).

24. Smirnov, K. & Sitnikova, E. Developmental milestones and behavior of infant rats: The role of sensory input from whiskers. Behavioural Brain Research 374, 112143 (2019).

25. Fuochi, S. et al. Puberty onset curve in CD (Sprague Dawley) and Long Evans outbred male rats. Lab. Anim. 56, 471–475 (2022).

26. Asiminas, A. et al. Experience-dependent changes in hippocampal spatial activity and hippocampal circuit function are disrupted in a rat model of Fragile X Syndrome. Mol. Autism 13, 49 (2022).

27. Ivanova, D. et al. Control of synaptic vesicle release probability via VAMP4 targeting to endolysosomes. Sci. Adv. 7, 1–22 (2021).

28. Bonnycastle, K. et al. Reversal of cell, circuit and seizure phenotypes in a mouse model of DNM1 epileptic encephalopathy. Nat. Commun. 14, 5285 (2023).

29. Subrahmanyam, R. et al. Reciprocal regulation of spontaneous synaptic vesicle fusion by Fragile X mental retardation protein and group I metabotropic glutamate receptors. J. Neurochem. 158, 1094–1109 (2021).

30. Kim, S. H. & Ryan, T. A. CDK5 Serves as a Major Control Point in Neurotransmitter Release. Neuron 67, 797–809 (2010).

31. Kim, S. H. & Ryan, T. A. CDK5 Serves as a Major Control Point in Neurotransmitter Release. Neuron 67, 797–809 (2010).

32. Dirkx, N., Miceli, F., Taglialatela, M. & Weckhuysen, S. The Role of Kv7.2 in Neurodevelopment: Insights and Gaps in Our Understanding. Front. Physiol. 11, (2020).

33. Rodriguez Gotor, J. J., Mahfooz, K., Perez-Otano, I. & Wesseling, J. F. Parallel processing of quickly and slowly mobilized reserve vesicles in hippocampal synapses. Elife 12, (2024).

34. Contractor, A., Klyachko, V. A. & Portera-Cailliau, C. Altered Neuronal and Circuit Excitability in Fragile X Syndrome. Neuron 87, 699–715 (2015).

35. Myrick, L. K. et al. Independent role for presynaptic FMRP revealed by an *FMR1* missense mutation associated with intellectual disability and seizures. Proceedings of the National Academy of Sciences 112, 949–956 (2015).

36. Shah, M. M., Migliore, M., Valencia, I., Cooper, E. C. & Brown, D. A. Functional significance of axonal Kv7 channels in hippocampal pyramidal neurons. Proceedings of the National Academy of Sciences 105, 7869–7874 (2008).

37. Gordon, S. L. & Cousin, M. A. X-Linked Intellectual Disability-Associated Mutations in Synaptophysin Disrupt Synaptobrevin II Retrieval. The Journal of Neuroscience 33, 13695–13700 (2013).

38. Gordon, S. L., Harper, C. B., Smillie, K. J. & Cousin, M. A. A Fine Balance of Synaptophysin Levels Underlies Efficient Retrieval of Synaptobrevin II to Synaptic Vesicles. PLoS One 11, e0149457 (2016).

39. Lai, H. C. & Jan, L. Y. The distribution and targeting of neuronal voltage-gated ion channels. Nat. Rev. Neurosci. 7, 548–562 (2006).

40. Gutzmann, A. et al. A period of structural plasticity at the axon initial segment in developing visual cortex. Front. Neuroanat. 8, (2014).

41. Bolós, M. et al. Maturation dynamics of the axon initial segment (AIS) of newborn dentate granule cells in young adult C57BL/6J mice. The Journal of Neuroscience 2253–18 (2019) doi:10.1523/JNEUROSCI.2253-18.2019.

42. Jamann, N. et al. Sensory input drives rapid homeostatic scaling of the axon initial segment in mouse barrel cortex. Nat. Commun. 12, 23 (2021).

43. Fujitani, M., Otani, Y. & Miyajima, H. Pathophysiological Roles of Abnormal Axon Initial Segments in Neurodevelopmental Disorders. Cells 10, 2110 (2021).

44. Kroon, T., van Hugte, E., van Linge, L., Mansvelder, H. D. & Meredith, R. M. Early postnatal development of pyramidal neurons across layers of the mouse medial prefrontal cortex. Sci. Rep. 9, 5037 (2019).

45. Hunsberger, M. S. & Mynlieff, M. BK potassium currents contribute differently to action potential waveform and firing rate as rat hippocampal neurons mature in the first postnatal week. J. Neurophysiol. 124, 703–714 (2020).

46. Dovzhenok, A. & Kuznetsov, A. S. Exploring Neuronal Bistability at the Depolarization Block. PLoS One 7, e42811 (2012).

47. Shao, L., Halvorsrud, R., Borg-Graham, L. & Storm, J. F. The role of BK-type Ca ^2+^-dependent K ^+^ channels in spike broadening during repetitive firing in rat hippocampal pyramidal cells. J. Physiol. 521, 135–146 (1999).

48. Leterrier, C. The Axon Initial Segment: An Updated Viewpoint. The Journal of Neuroscience 38, 2135–2145 (2018).

49. Yamada, R. & Kuba, H. Structural and Functional Plasticity at the Axon Initial Segment. Front. Cell. Neurosci. 10, (2016).

50. Booker, S. A. et al. Input-Output Relationship of CA1 Pyramidal Neurons Reveals Intact Homeostatic Mechanisms in a Mouse Model of Fragile X Syndrome. Cell Rep. 32, 107988 (2020).

51. Rosa, F. et al. KV7.2 channel dysfunction delays neuronal maturation and undermines early network development in a hiPSC model of KCNQ2-DEE. Neurobiol. Dis. 216, 107120 (2025).

52. Asiminas, A. et al. Sustained correction of associative learning deficits following brief, early treatment in a rat model of Fragile X Syndrome. Sci. Transl. Med. 11, 1–11 (2019).

53. Jain, A. et al. Presynaptic filopodia form kinapses and modulate membrane mechanics for synchronous neurotransmission and seizure generation. bioRxiv (2024) doi:10.1101/2024.10.07.616970.

54. Hönigsperger, C., Marosi, M., Murphy, R. & Storm, J. F. Dorsoventral differences in Kv7/M-current and its impact on resonance, temporal summation and excitability in rat hippocampal pyramidal cells. J. Physiol. 593, 1551–1580 (2015).

