## Supplementary Information for "Altered axonal initial segment development links circuit and K_v_7 dysfunction in an *Fmr1* knockout rat"

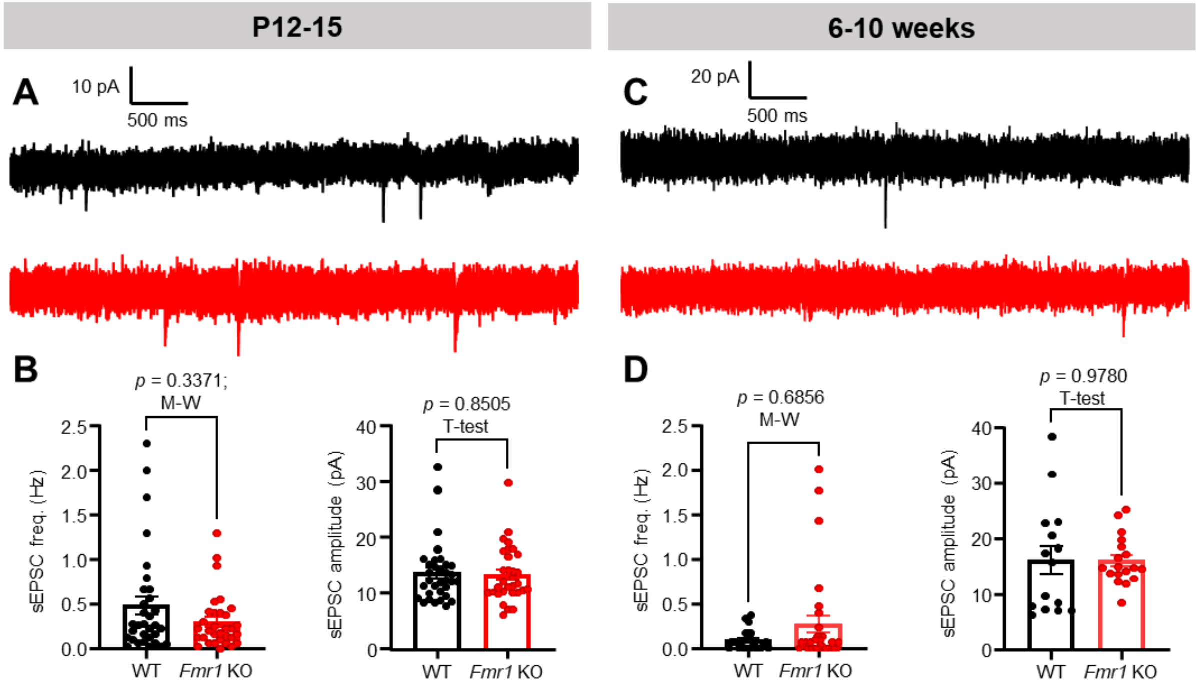


**Figure S1: Spontaneous excitatory synaptic drive onto CA1 pyramidal neurons in male *Fmr1* KO rats is unaffected throughout development. (A)** Example traces of spontaneous excitatory postsynaptic currents (sEPSCs) recorded in CA1 pyramidal neurons in acute slices from P12-15 *Fmr1* KO vs WT rats. **(B)** Comparison plots of sEPSC frequency and amplitude in both conditions. Frequency = 32/11 WT vs 32/13 KO neurons / rats; amplitude = 32/11 WT vs 32/13 KO neurons / rats. **(C)** and **(D)** As for (**A**) and (**B**), in 6-10-week old rats. Frequency = 24/11 WT vs 30/8 KO; amplitude = 15/11 WT vs 18/13 KO. All summary data are mean ± SEM.

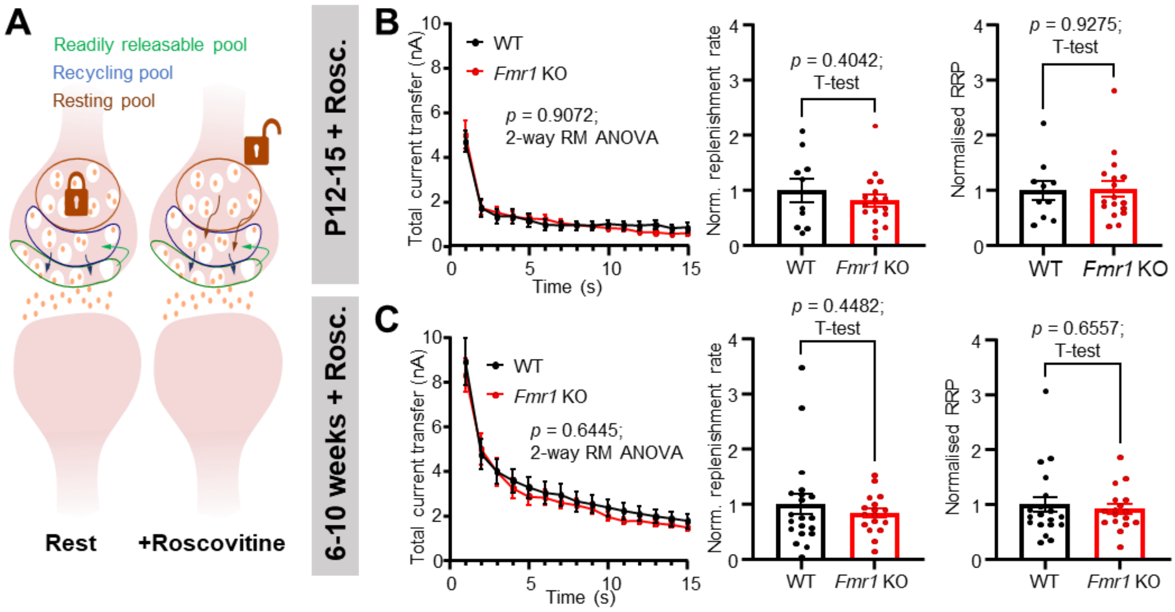


**Figure S2. *Fmr1* KO does not alter the static size of the RRP: (A)** Schematic showing mechanism of action of the Cdk5 inhibitor roscovitine. Roscovitine unlocks access to the resting vesicle pool, thus allowing a greater number of vesicles to be recruited during neuronal activity. (**B**) Left: Plot of total current transfer binned per second during a presynaptic SC 40 Hz stimulation train of 600 action potentials recorded in CA1 pyramidal neurons from slices pretreated with 100 µM roscovitine (Rosc.) in P12-15 *Fmr1* KO vs WT rats. Middle: Comparison of vesicle replenishment rates and right; Readily Releasable Pool (RRP) estimates from the same experiments. 10/5 WT vs 17/9 KO for all comparisons neurons / rats. (**C**) As for (**B**), in 6–10-week-old rats. 20/8 WT vs 17/6 for all comparisons neurons / rats. All summary data are mean ± SEM. n/N = neurons / rats.

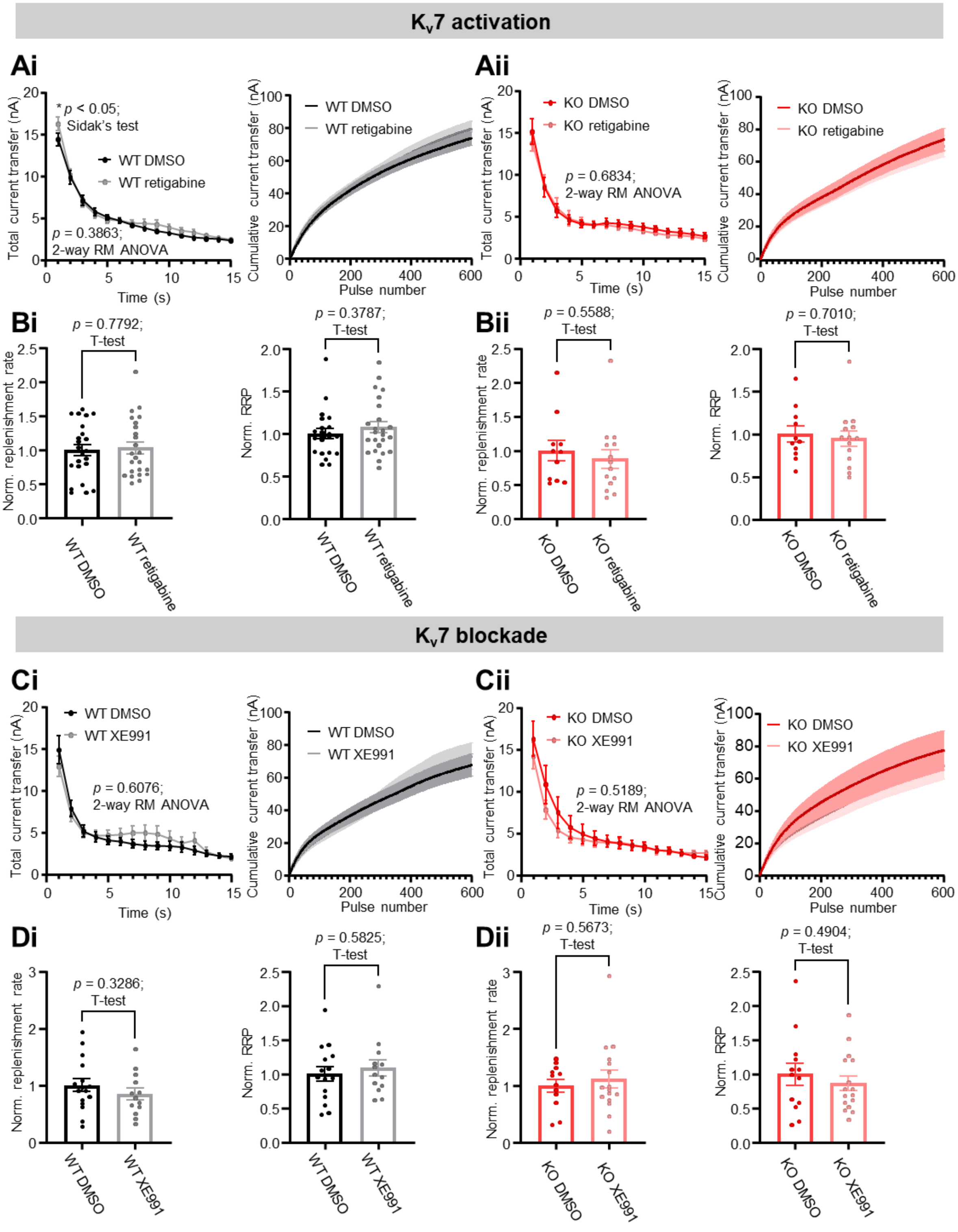


**Figure S3. Effect of K$_v$7 channel modulation on SV pool replenishment in P12-15 *Fmr1* KO rats. (Ai)** Left: Total current transfer measured during a presynaptic 40Hz stimulation train of 600 action potentials binned as total current per second and right: cumulative current transfer over the course of stimulation recorded in acute slices from WT rats pretreated with either 0.1 % DMSO or 2.5 µM retigabine. 23/13 DMSO vs 24/13 retigabine neurons / rats. **(Bi)** Left: comparison of vesicle replenishment rates and right: Readily Releasable Pool (RRP) estimates extrapolated from graphs in (**Ai**). Details on calculations for these measurements can be found in methods. 23/13 DMSO vs 24/13 retigabine neurons / rats. **(Aii & Bii)** As for

(**Ai**) and (**Bi**) in *Fmr1* KO rats. **(Aii)** 11/7 DMSO vs 14/7 retigabine neurons / rats. (**Bii**) 11/7 DMSO vs 14/7 retigabine neurons / rats. **(C-D)** As for (**A-B**), treated with 5 μM XE991 or 0.1 % DMSO. WT = 16/10 DMSO vs 13/9 XE991; KO = 13/10 DMSO vs 13/9 XE neurons / rats. All summary data are mean ± SEM. n/N = neurons / rats.

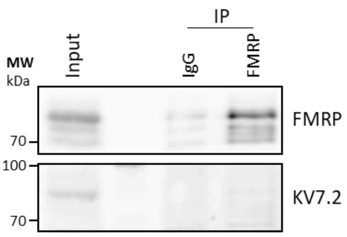


**Figure S4. FMRP does not directly interact with $K_v$7.2**. Western blot of immunoprecipitation assay showing a lack of interaction between FMRP and $K_v$7.2 in P14 WT rat brain lysates. Either FMRP or control IgG immunoprecipitates were probed for both FMRP and $K_v$7.2. Immunoprecipitate input is also displayed (10 % of total). Western blots are representative of experiments with N = 3 rats.

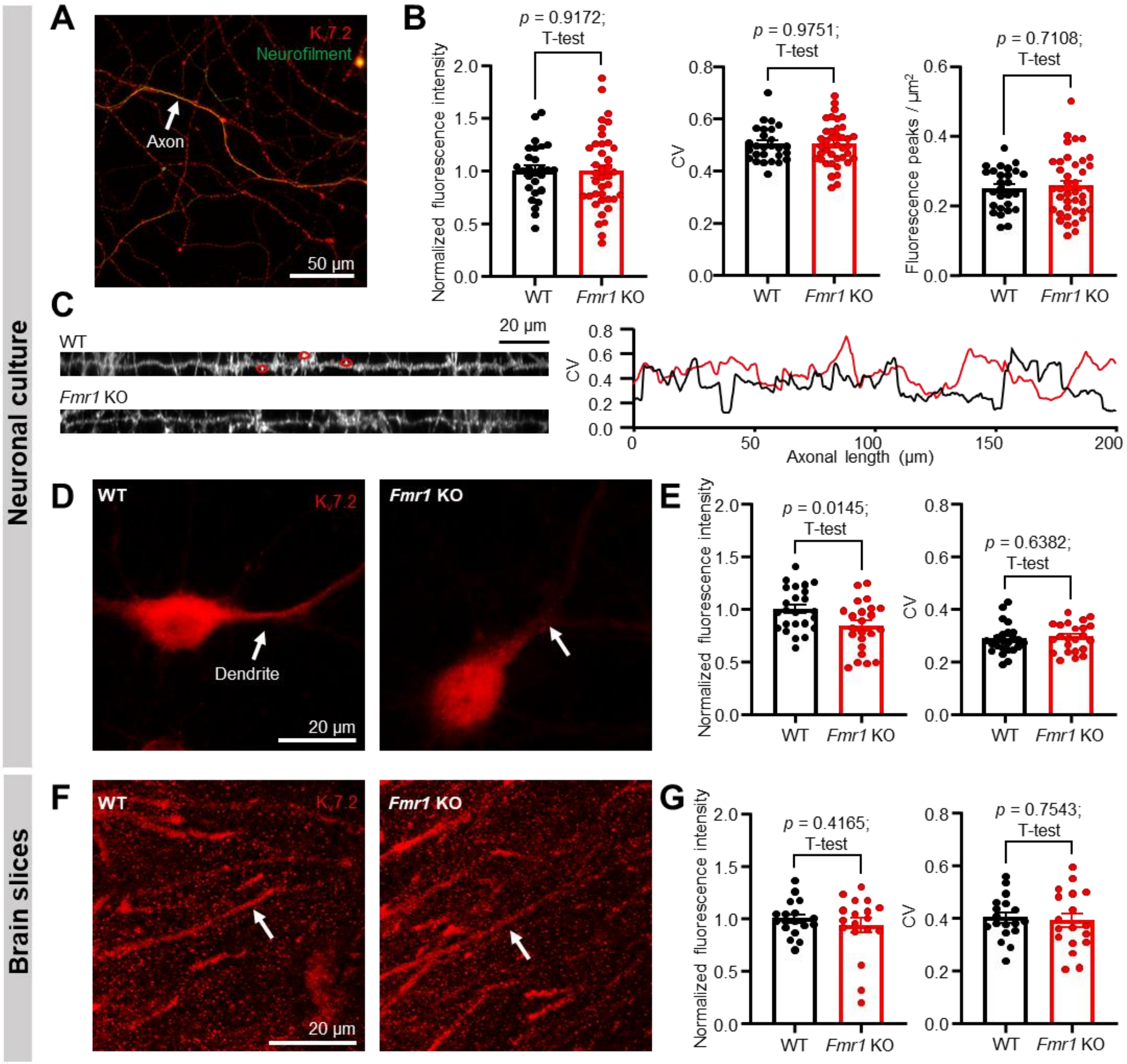


**Figure S5: K$_v$7.2 gross expression and clustering is broadly unaltered in *Fmr1* KO rat neurites. (A)** Example images depicting the K$_v$7.2 staining profiles along the axon of cultured WT hippocampal neurons. **(B)** Comparison plots of normalised fluorescence intensity, CV, and detected fluorescence peaks in axons of WT vs *Fmr1* KO neurons.**(C)** Left: example outputs of background-corrected axonal K$_v$7.2 intensity profiles in WT vs *Fmr1* KO neurons. Red circles indicate fluorescence peaks. Right: example coefficient of variation (CV) profile comparison derived from the two above images; an indication of channel clustering. **(D)** Example images of K$_v$7.2 staining profiles in the dendrites of cultured WT vs *Fmr1* KO neurons. **(E)** Comparison plots of normalised fluorescence intensity and CV in dendrites of WT vs *Fmr1* KO neurons. Arrows indicate portions of dendrites analysed. 27/7 WT vs 39/11 KO coverslips / embryos for all primary neuronal culture comparisons. **(F)** and (**G**) As for (**D**) and (**E**), in fixed brain slices. 18/6 WT vs 18/6 KO slices / rats for all brain slice comparisons.

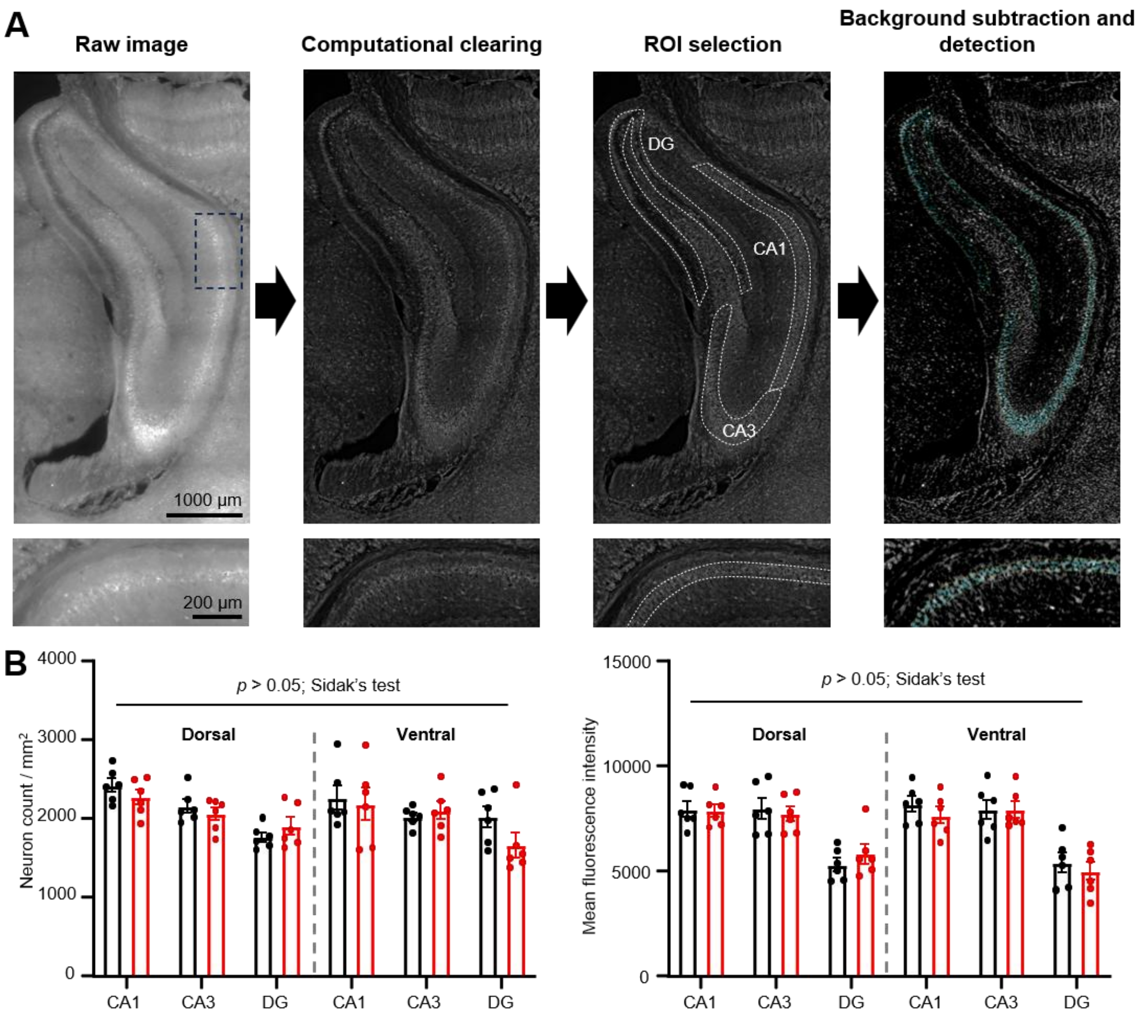


**Figure S6: Automated regional detection of $K_v7.2$-positive neurons. (A)** Top: example images of $K_v7.2$ expression throughout the hippocampal formation outlining workflow. Raw tile scans (left) were computationally cleared using THUNDER software (middle left). Regions of interest (ROI's) are annotated (middle right), and then custom software performs background subtraction on the image, and automatically detects $K_v7.2$-positive neurons within each ROI (right; cyan markers depict detected neurons). Bottom: zoom insets of the boxed CA1 region highlighted on the top leftmost image. **(B)** Quantification of total positive neuron counts (left) and mean fluorescence intensity of detected neurons (right) in each hippocampal sub-region in *Fmr1* KO slices vs WT littermate controls. CA = cornu ammonis; DG = dentate gyrus. N = 6 rats for both WT and KO conditions.

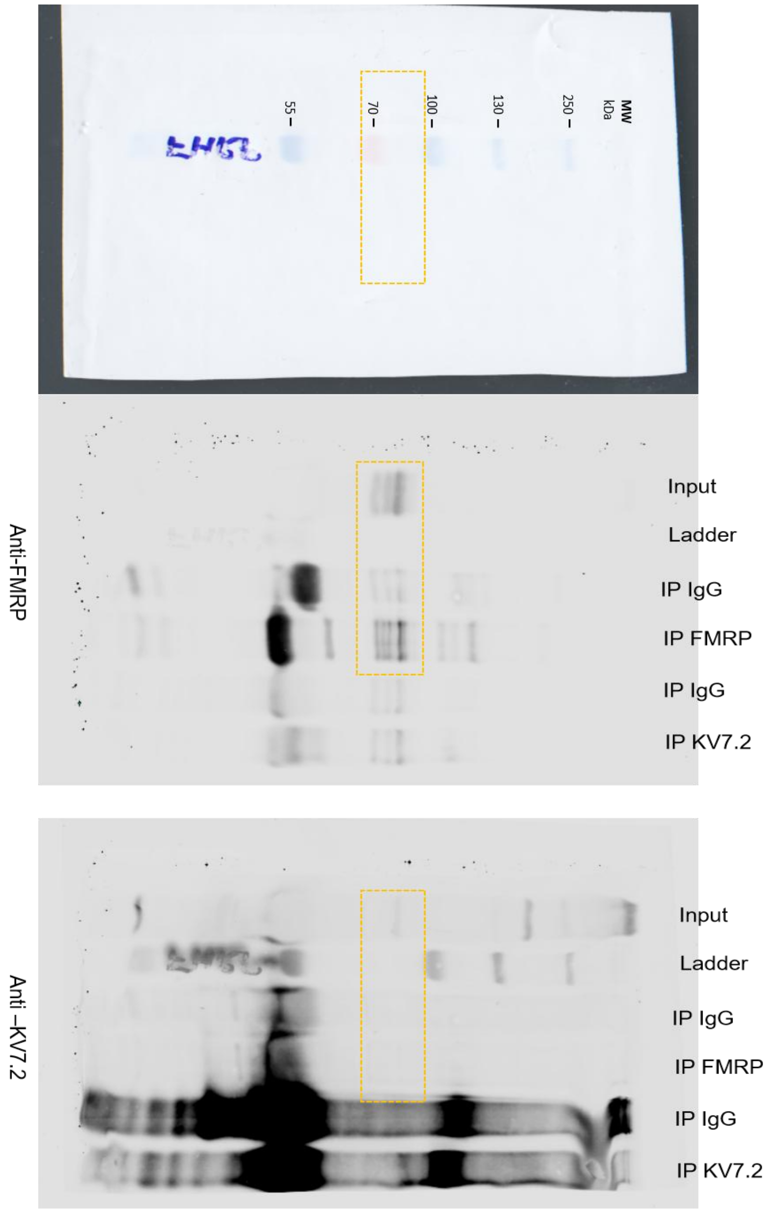


**Figure S7: Raw uncropped immunoprecipitation blots used to create Figure S4.**

| | WT | | KO | | | |
|---|---|---|---|---|---|---|
| Property | Mean / median | SEM (±) | Mean / median | SEM (±) | *p* value | Statistical test |
| RMP (mV) | -60.35 | 0.82 | -62.31 | 0.75 | 0.0846 | T-test |
| $R_{in}$ (MΩ) | 159.50 | 6.66 | 170.70 | 7.99 | 0.2883 | T-test |
| Rheobase (pA) | 69.85 | 3.91 | 68.57 | 3.45 | 0.8066 | T-test |
| Sag (mV) | 19.67 | 0.75 | 21.55 | 0.70 | 0.0728 | T-test |
| AP threshold (mV) | -45.30 | 0.71 | -43.53 | 0.96 | 0.1483 | T-test |
| AP amplitude (mV) | 107.40 | 0.93 | 105.10 | 1.25 | 0.1495 | T-test |
| AP RT (ms) | 0.37 | 0.27 | 0.38 | 1.25 | 0.5844 | T-test |
| AP HW (ms) | 1.99 | 0.04 | 2.04 | 0.05 | 0.5070 | T-test |
| AP latency (ms) | 71.15 | 9.57 | 83.05 | 15.74 | 0.0600 | Mann-Whitney |
| fAHP (mV) | 11.25 | 0.4456 | 10.99 | 0.5722 | 0.8534 | Mann-Whitney |
| mAHP (mV) | 12.74 | 0.5083 | 13.24 | 0.5784 | 0.5186 | T-test |

**Table S1: Intrinsic properties of CA pyramidal neurons recorded acute slices from P12-15 *Fmr1* KO vs WT rats.** Note that mean value is given where a T-test was used, and median value is given where a Mann-Whitney test was used (methods). All action potential property values are derived from the rheobase action potential. Abbreviations: RMP = resting membrane potential; $R_{in}$ = input resistance; AP = action potential; RT = rise time; HW = half-width; fAHP = fast afterhyperpolarisation; mAHP = medium afterhyperpolarisation.

| | WT | | KO | | | |
|---|---|---|---|---|---|---|
| Property | Mean / median | SEM (±) | Mean / median | SEM (±) | *p* value | Statistical test |
| RMP (mV) | -67.05 | 1.09 | -64.65 | 1.26 | 0.1566 | Mann-Whitney |
| $R_{in}$ (MΩ) | 74.30 | 3.95 | 82.59 | 6.88 | 0.0940 | Mann-Whitney |
| Rheobase (pA) | 125.00 | 9.46 | 125.00 | 6.85 | 0.3282 | Mann-Whitney |
| Sag (mV) | 10.32 | 0.65 | 10.31 | 0.73 | 0.9908 | T-test |
| AP threshold (mV) | -44.27 | 0.76 | -45.25 | 0.52 | 0.2967 | T-test |
| AP amplitude (mV) | 107.70 | 0.96 | 107.90 | 1.45 | 0.8926 | Mann-Whitney |
| AP RT (ms) | 0.36 | 0.02 | 0.35 | 0.01 | 0.6837 | T-test |
| AP HW (ms) | 2.01 | 0.03 | 2.01 | 0.04 | 0.9355 | T-test |
| AP latency (ms) | 50.45 | 10.00 | 51.70 | 17.29 | 0.5966 | Mann-Whitney |
| fAHP (mV) | 18.77 | 0.5042 | 18.07 | 0.5658 | 0.3710 | T-test |
| mAHP (mV) | 16.94 | 0.4972 | 16.48 | 0.8094 | 0.6327 | T-test |

**Table S2: Intrinsic properties of CA pyramidal neurons recorded acute slices from 6 to 10-week old *Fmr1* KO vs WT rats.** Note that mean value is given where a T-test was used, and median value is given where a Mann-Whitney test was used (methods). All action potential property values are derived from the rheobase action potential. Abbreviations: RMP = resting membrane potential; $R_{in}$ = input resistance; AP = action potential; RT = rise time; HW = half-width: fAHP = fast afterhyperpolarisation; mAHP = medium afterhyperpolarisation.

| | WT DMSO | | WT Retigabine | | | |
|---|---|---|---|---|---|---|
| Property | Mean / median | SEM (±) | Mean / median | SEM (±) | *p* value | Statistical test |
| RMP (mV) | -60.61 | 0.6107 | -60.26 | 0.8753 | 0.7376 | T-test |
| $R_{in}$ (MΩ) | 168.3 | 4.591 | 162.3 | 6.482 | 0.4532 | T-test |
| AP threshold (mV) | -43.95 | 0.9276 | -43.97 | 0.6797 | 0.9851 | T-test |
| Rheobase fAHP (mV) | 12.61 | 0.4285 | 12.80 | 0.4599 | 0.7624 | T-test |
| Rheobase mAHP (mV) | 13.92 | 0.5114 | 13.92 | 0.4638 | 0.9899 | T-test |
| SFA index slope | 0.0577 | 0.0093 | 0.0721 | 0.0100 | 0.2974 | T-test |

**Table S3: Intrinsic properties of CA1 pyramidal neurons recorded acute slices from P12-15 WT rats pretreated with either 0.1 % DMSO or 2.5 µM retigabine.** Note that mean value is given where a T-test was used, and median value is given where a Mann-Whitney test was used (methods). Abbreviations: RMP = resting membrane potential; $R_{in}$ = input resistance; AP = action potential; fAHP = fast afterhyperpolarisation; mAHP = medium afterhyperpolarisation; SFA = spike frequency adaptation.

| | KO DMSO | | KO Retigabine | | | |
|---|---|---|---|---|---|---|
| Property | Mean / median | SEM (±) | Mean / median | SEM (±) | *p* value | Statistical test |
| RMP (mV) | -59.59 | 0.9081 | -60.49 | 0.6047 | 0.3965 | T-test |
| $R_{in}$ (MΩ) | 186.1 | 14.38 | 190.5 | 11.02 | 0.6565 | Mann-Whitney |
| AP threshold (mV) | -45.67 | 0.8878 | -45.75 | 0.8922 | 0.4925 | Mann-Whitney |
| Rheobase fAHP (mV) | 11.22 | 0.5427 | 11.88 | 0.5221 | 0.4708 | Mann-Whitney |
| Rheobase mAHP (mV) | 13.39 | 0.5944 | 13.60 | 0.5467 | 0.7950 | T-test |
| SFA index slope | 0.05181 | 0.0100 | 0.05737 | 0.0093 | 0.3667 | Mann-Whitney |

**Table S4: Intrinsic properties of CA1 pyramidal neurons recorded acute slices from P12-15 *Fmr1* KO rats pretreated with either 0.1 % DMSO or 2.5 µM retigabine.** Note that mean value is given where a T-test was used, and median value is given where a Mann-Whitney test was used (methods). Abbreviations: RMP = resting membrane potential; $R_{in}$ = input resistance; AP = action potential; fAHP = fast afterhyperpolarisation; mAHP = medium afterhyperpolarisation; SFA = spike frequency adaptation.

| | WT DMSO | | WT XE991 | | | |
|---|---|---|---|---|---|---|
| Property | Mean / median | SEM (±) | Mean / median | SEM (±) | *p* value | Statistical test |
| RMP (mV) | -63.55 | 0.8854 | -58.30 | 1.2590 | 0.0011 | T-test |
| $R_{in}$ (MΩ) | 147.6 | 6.342 | 157.8 | 7.465 | 0.2986 | T-test |
| AP threshold (mV) | -46.09 | 0.7114 | -44.77 | 0.9300 | 0.2605 | T-test |
| Rheobase fAHP (mV) | 10.71 | 0.4905 | 10.49 | 0.5637 | 0.7610 | Mann-Whitney |
| Rheobase mAHP (mV) | 12.93 | 0.6365 | 12.57 | 0.5114 | 0.5807 | Mann-Whitney |
| SFA index slope | 0.0665 | 0.0207 | 0.0609 | 0.0185 | 0.8436 | T-test |

**Table S5: Intrinsic properties of CA1 pyramidal neurons recorded acute slices from P12-15 WT rats pretreated with either 0.1 % DMSO or 5 µM XE991.** Note that mean value is given where a T-test was used, and median value is given where a Mann-Whitney test was used (methods). Abbreviations: RMP = resting membrane potential; $R_{in}$ = input resistance; AP = action potential; fAHP = fast afterhyperpolarisation; mAHP = medium afterhyperpolarisation; SFA = spike frequency adaptation.

| | KO DMSO | | KO XE991 | | | |
|---|---|---|---|---|---|---|
| **Property** | **Mean / median** | **SEM (±)** | **Mean / median** | **SEM (±)** | ***p* value** | **Statistical test** |
| RMP (mV) | -59.56 | 0.9062 | -58.96 | 0.9130 | 0.6513 | T-test |
| $R_{in}$ (MΩ) | 152.4 | 6.485 | 153.7 | 4.050 | 0.8650 | T-test |
| AP threshold (mV) | -45.65 | 0.7031 | -44.43 | 0.7579 | 0.2460 | T-test |
| Rheobase fAHP (mV) | 12.00 | 0.4457 | 10.31 | 0.5496 | 0.0202 | T-test |
| Rheobase mAHP (mV) | 13.54 | 0.4522 | 12.46 | 0.5350 | 0.1280 | T-test |
| SFA index slope | 0.0609 | 0.0149 | 0.0610 | 0.0164 | 0.9960 | T-test |

**Table S6: Intrinsic properties of CA1 pyramidal neurons recorded acute slices from P12-15 *Fmr1* KO rats pretreated with either 0.1 % DMSO or 5 µM XE991.** Note that mean value is given where a T-test was used, and median value is given where a Mann-Whitney test was used (methods). Abbreviations: RMP = resting membrane potential; $R_{in}$ = input resistance; AP = action potential; fAHP = fast afterhyperpolarisation; mAHP = medium afterhyperpolarisation; SFA = spike frequency adaptation.
